# Intrinsic molecular susceptibility underlies selective neuronal vulnerability in the Alzheimer’s disease entorhinal cortex

**DOI:** 10.64898/2026.06.22.733753

**Authors:** Samuel L Boulger, Daniela Lerose, Emily Adair, Marianna Papageorgopoulou, Yinuo Zhao, Baptiste Avot, Michael Thomas, Pavit Nibber, Hannah Klug, Nurun N Fancy, Anna Mallach, Eugene P Duff, Alessia Caramello, Paul M Matthews

## Abstract

Entorhinal cortex (EC) excitatory neurons are lost early in Alzheimer’s disease (AD), yet the specific subtype and characteristics contributing to this vulnerability are poorly understood. Combining imaging mass cytometry (206,913 cells; 62 donors) and single nucleus RNA sequencing (42,780 nuclei; 36 donors) of *post-mortem* EC, we found that calbindin-expressing layer 2-3 excitatory neurons accumulate high phospho-tau burden and are preferentially lost in AD. In non-diseased brains, these neurons exhibit elevated tau-modifying kinase expression (ERK1/2, FYN, ROCK), reduced phosphatase expression (PP2A/B, PP5) and low mitochondrial respiratory capacity which together are predicted to promote high vulnerability to tau pathology. Trajectory analysis resolved progression from homeostasis through DNA damage and proteostatic stress to developmental re-entry and death priming. *In silico* screening suggested histone deacetylase inhibitors and cyclooxygenase inhibitors as candidate resilience-promoting therapeutics. Our work thus reframes intrinsic features of neuronal identity promoting phospho-tau formation as modifiable determinants of the selective vulnerability of EC calbindin neurons.

## Introduction

While neuronal and synaptic pathology is widespread in brains of patients with Alzheimer’s disease (AD)^1^, substantial neuronal loss is strikingly selective^2^. The pathological hallmarks of disease – amyloid-β (Aβ) plaques^3^ and tau neurofibrillary tangles^4^ – do not accumulate uniformly, and progress in a consistent, region-specific fashion over time. In affected regions, specific neuronal populations are severely affected while anatomically adjacent cells are largely spared^5^. Understanding the cellular and molecular basis of this selectivity is a fundamental question in AD biology.

It has long been established that excitatory neurons in layer 2 of the entorhinal cortex (EC) are amongst the first to die in AD^6–8^. Beyond that, there are limited molecular descriptions of vulnerable EC neurons in human AD brains, but studies have indicated that calbindin-^9–11^, reelin-^12^, and RORβ-expressing^13^ excitatory neurons, and parvalbumin^9,14^, somatostatin^15,16^, and neuropeptide Y-expressing^16,17^ interneurons are particularly susceptible to loss as pathology progresses.

Rodent models of AD have provided mechanistic insights into subtype-specific vulnerability: EC reelin-positive neurons have been shown in rats to accumulate Aβ intracellularly as a consequence of elevated *App* expression driven by their large size and metabolic demands^18,19^, and we have previously shown that intracellular Aβ accumulation associates with neuronal loss in the human middle temporal gyrus in AD^20^. However, the molecular features that define EC neurons in the human brain, and whether these represent intrinsic properties of neuronal identity or consequences of pathological exposure, remain unresolved.

Here, we sought to provide a comprehensive molecular characterisation of selectively vulnerable neurons in the human EC, and to test the hypothesis that vulnerability reflects an intrinsic property of neuronal identity rather than simply arising as a consequence of exposure to pathology. Using imaging mass cytometry (IMC) and single nucleus RNA sequencing (snRNA-seq), we identified neuronal populations that are preferentially lost with AD progression and profiled transcriptomic differences between vulnerable and resilient populations to infer the molecular pathways underpinning vulnerability. By integrating proteomic and transcriptomic readouts across disease severity, we defined the pathological trajectory of vulnerable neurons from a homeostatic state through progressive dysfunction to cell death. Together, these findings identify intrinsic molecular features of vulnerable neurons that are independent of overt pathology and predict downstream tau burden, supporting a model in which selective vulnerability reflects neuronal identity as well as exposure to pathology stress. We further leverage the resulting resilience signature in an *in silico* drug repurposing analysis to nominate candidate small molecules aligned with the resilient transcriptional state, providing a mechanistic and translational framework for therapeutic strategies to enhance resilience in the human EC.

## Results

### Identification of entorhinal neuronal subtypes and their cortical organisation using parallel imaging mass cytometry and single nucleus transcriptomics

IMC and snRNA-seq were performed on *post-mortem* EC from individuals with and without AD pathology (Fig. 1a, 1b; Supplementary Table 1). IMC was first performed to identify neuronal subtypes lost with AD pathology using an optimised panel of 34 antibodies enriched for neuronal markers, along with a DNA intercalator (Supplementary Fig. 1). In the same antibody panel, markers for AD pathology and protein degradation were also included for consecutive exploration of disease mechanisms. Following image preprocessing and quality control, 215,739 cells were identified across 67 donors (1106 ± 300 cells per region of interest; 916 ± 243 cells per mm^2^). Analyses were performed only on samples retaining all three ROIs after quality control (n = 18 non-diseased donors (Braak 0-1; 52,929 cells); n = 19 mid-stage AD (Braak 3-4; 62,320 cells); n = 25 late-stage AD (Braak 6; 91,664 cells)). The OLIG2 marker was excluded due to suboptimal performance.

**Fig. 1.**
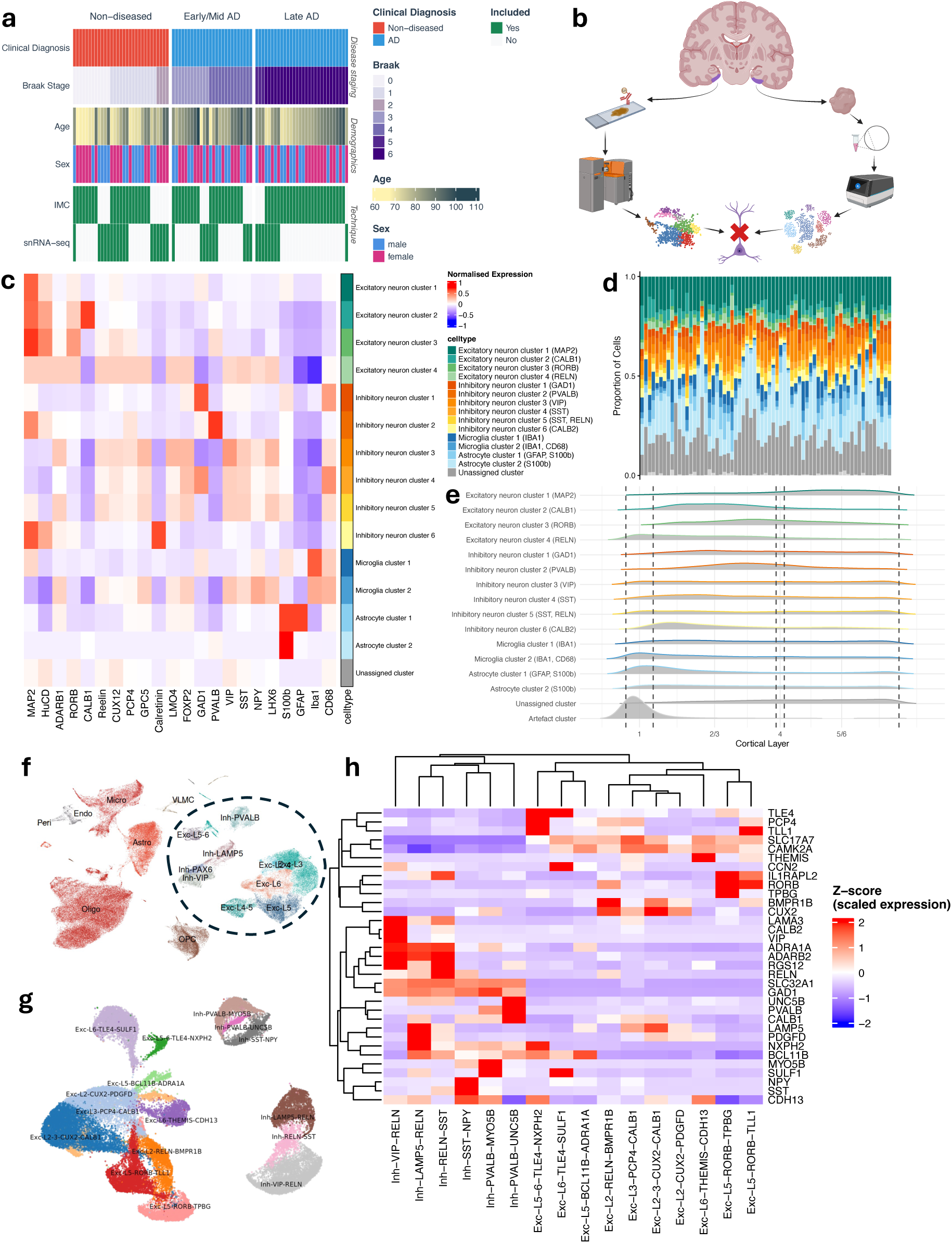
Cell type assignment of nuclei identified by IMC and snRNA-seq in the human entorhinal cortex. a, Clinicopathological metadata for the IMC and snRNA-seq cohorts, stratified by disease stage group. **b,** Schematic of the parallel IMC and snRNA-seq workflow applied to human post-mortem EC. Created in BioRender. Boulger, S. (2026) https://BioRender.com/q1qqafd. **c,** Heatmap of normalised expression of lineage and cell type-defining markers across major IMC-identified cell types; markers shown in the heatmap are those used for clustering and annotation (type markers). **d,** Stacked bar plots showing relative proportional abundance of annotated cell types among all cells passing quality control, stratified by donor. **e,** Spatial density maps depicting the laminar distribution of annotated cell types across entorhinal cortex layers. Layer boundaries were approximated from published histological atlases^24^. **f,** UMAP projection of initial snRNA-seq clusters. **g,** UMAP projection of snRNA-seq neuronal populations following two rounds of subclustering. **h,** Heatmap of normalised expression of entorhinal cortex neuronal subtype-defining markers across neuronal subclusters^23^.

Cells were clustered using a two-step unsupervised approach and assigned to neuronal or glial subpopulations based on cell-type-specific marker expression (Fig. 1c; Supplementary Fig. 2; Supplementary Table 2). Of the clustered cells, 24.2 ± 5.7% (mean ± standard deviation) were classified as excitatory neurons, and 24.5 ± 9.8% as inhibitory neurons (Fig. 1d). 8.6 ± 5.3% were microglia, 23.6 ± 7.7% were astrocytes, and 19.2 ± 9.2% were left unassigned (Fig. 1d). 14 neuronal and glial subpopulations were identified based on distinct marker profiles (Fig. 1c), and their spatial distribution revealed laminar organisation consistent with known EC architecture^21^ (Fig. 1e).

For snRNA-seq, nuclei were isolated and RNA sequenced from frozen tissue from a subset (n = 19) of the full IMC cohort alongside 2 additional mid-stage AD samples. These data were combined with a cohort of EC snRNA-seq data from a further 19 samples previously described^22^. 4 samples were excluded during quality control assessment, leaving a cohort of non-diseased controls (n = 17 donors, 21,884 nuclei; Braak 0-2; no AD diagnosis), individuals with mid-stage AD (n = 10 donors, 9622 nuclei; Braak 3-4; AD diagnosis), and late-stage AD (n = 9 donors, 11,274 nuclei; Braak 5-6; AD diagnosis). After initial clustering (Fig. 1f) and two rounds of neuronal subclustering (Fig. 1g), we identified 10 excitatory and 6 inhibitory neuronal populations (Supplementary Table 3), which were annotated based primarily on previously defined entorhinal cortex neuronal subtype-specific marker genes^23^ (Fig. 1h; Supplementary Table 4).

### Layer 2-3 calbindin-expressing excitatory neurons are selectively vulnerable to loss

To assess disease-associated changes in sizes of different cell populations, we quantified columnar density – the number of cells per mm along the cortical ribbon spanning the full grey matter depth. This metric normalises for variation in cortical thickness between samples and therefore distinguishes neuronal loss from apparent density changes arising from cortical atrophy^25,26^. Using IMC, we observed a selective reduction in excitatory neuron cluster 2 (CALB1), representing layer 2-3 calbindin-positive pyramidal neurons, in patients diagnosed with AD (Fig. 2a; Supplementary Table 5, 6). Separating donors by Braak stage group revealed that this loss is already apparent at mid-stage disease (Braak 3-4; Fig. 2b). Several inhibitory neuron populations were comparatively preserved, and excitatory clusters 3 and 4, representing RORβ and reelin-immunoreactive populations, trended towards a reduction with AD, but to a lesser degree than the calbindin-positive cluster and without reaching statistical significance (Fig. 2a). Simulation-based power was only 46% (excitatory cluster 3 (RORB)) and 12% (excitatory cluster 4 (RELN)) against a pre-specified threshold of log_2_FC = −0.5 (∼29% columnar density reduction; Supplementary Table 7); these clusters therefore cannot be considered preserved based on non-significance alone.

**Fig. 2.**
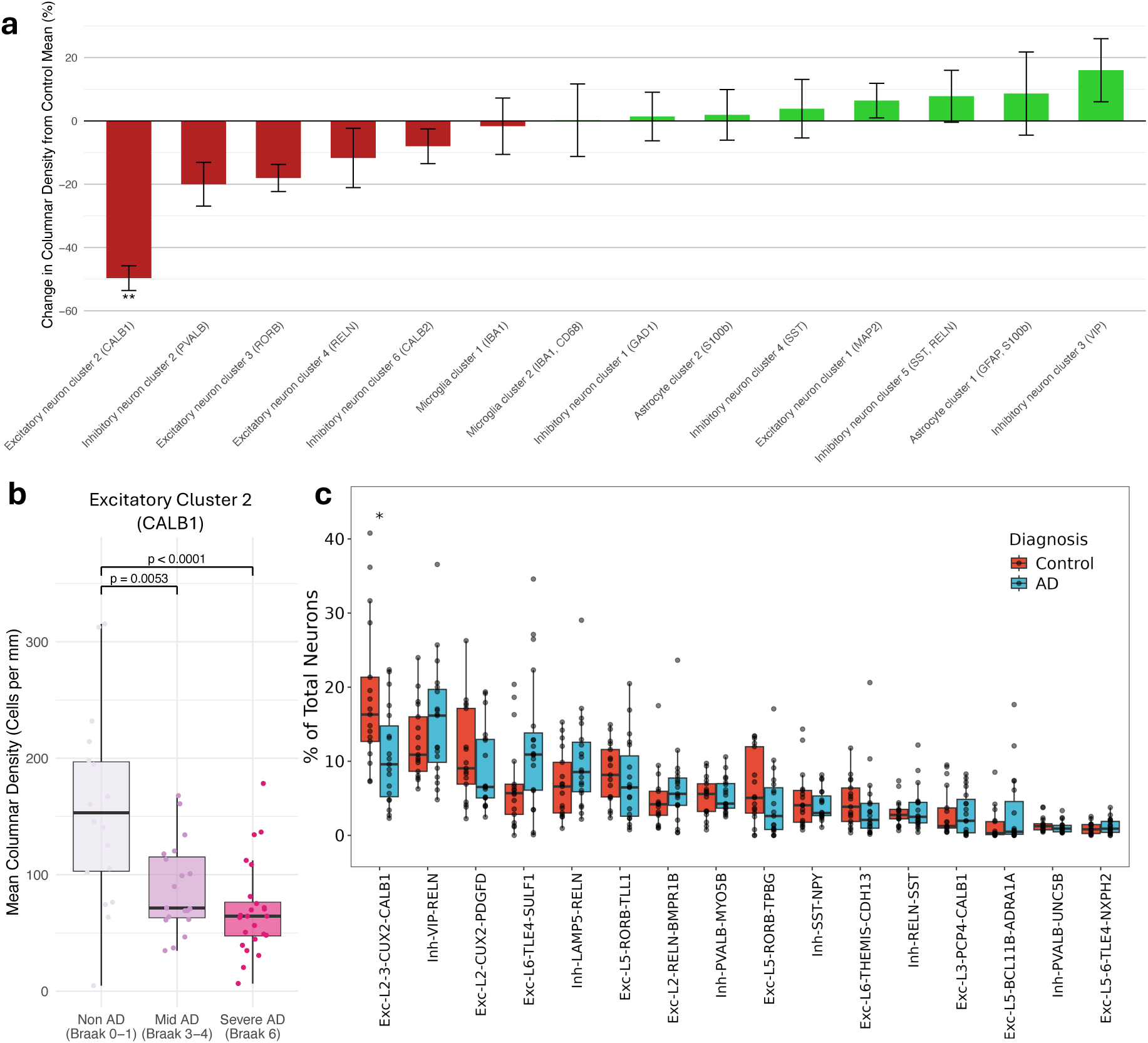
Selective loss of calbindin-expressing excitatory neurons in Alzheimer’s disease. a, Mean percentage change in columnar density relative to controls across all IMC-annotated cell types, ordered by magnitude of change. Columnar density was defined as the number of labelled cells per millimetre along the cortical column, measured orthogonally from pial surface to white matter, within each annotated region of interest. Error bars represent the SEM. Statistical comparisons were performed using linear mixed-effects models (columnar density ∼ diagnosis + age + (1|donor)); p values were adjusted using the Benjamini-Hochberg method. **b,** Columnar density of excitatory cluster 2 (CALB1) neurons stratified by Braak stage group (non-AD: Braak 0-1; mid AD: Braak 3-4; late-stage/severe AD: Braak 6). Statistical comparisons were performed using linear mixed-effects models (columnar density ∼ Braak group + age + (1|donor)); p values were adjusted for multiple comparisons using the Benjamini-Hochberg method within broad cell type groups. a-b, n = 18 non-diseased control donors, 19 mid AD, 25 late AD; IMC cohort with three ROIs passing quality control. **c,** Proportional abundance of snRNA-seq neuronal populations in AD versus control donors, ordered by population size. Statistics derived from Dirichlet multinomial regression (proportion ∼ diagnosis + age + sex + study ID). n = 17 control donors, 19 AD; snRNA-seq cohort. Each point represents one donor.

Consistent with these IMC findings, snRNA-seq independently revealed a selective reduction in *CALB1*-expressing excitatory neurons (Exc-L2-3-*CUX2*-*CALB1*; Fig. 2c; Supplementary Table 8), providing convergent evidence across orthogonal modalities for the selective vulnerability of this population. The mean relative proportion of Exc-L2-3-*CUX2*-*CALB1* was even lower when samples with earliest AD-related pathology (Braak 2) were excluded from the control group (Supplementary Fig. 3). The concordant loss of the transcriptomically defined Exc-L2-3-*CUX2*-*CALB1* population determined by snRNA-seq argues against downregulation of calbindin expression^10^ without neuronal loss as an explanation for the IMC findings.

To contextualise our findings within an existing large-scale transcriptomic atlas, we computed pairwise Jaccard similarity between our excitatory subcluster marker genes (Supplementary Table 3) and those reported in a previous publication showing neuronal subtype-specific loss in AD^12^. We revealed relatively high similarity between Exc-L2-3-*CUX2*-*CALB1* and the cluster Exc L2/3 *TOX3 TTC6* (Supplementary Fig. 4a), which was identified as significantly less abundant with AD, thereby validating our findings and extending previous work. We re-annotated our excitatory neurons using the published marker genes and observed statistically significant depletion of neurons corresponding to Exc *TOX3 TTC6* and Exc *AGBL1 GPC5* with Braak stage (Supplementary Fig. 4b). We did not observe the loss of Exc *RELN GPC5* or Exc *RELN COL5A2* neurons previously reported^12^.

### Layer 2-3 calbindin-expressing excitatory neurons are intrinsically vulnerable to tau pathology

Differential gene expression analysis, stratified by AD diagnosis, contrasting the Exc-L2-3-*CUX2*-*CALB1* population to all other excitatory neurons, revealed intrinsic features of these neurons that may contribute to their vulnerability in AD (Supplementary Table 9). Pathway enrichment revealed relative upregulation of postsynaptic glutamate receptor and calcium signalling pathways and downregulation of presynaptic vesicle exocytosis (Fig. 3a; Supplementary Table 10), a transcriptional signature consistent with the widespread synaptic inputs received by these large pyramidal neurons^27^ and with susceptibility to excitotoxic stress. Furthermore, pathway analysis uncovered altered transcriptional signatures related to proteostasis in Exc-L2-3-*CUX2*-*CALB1* neurons, including relatively reduced expression of Cul5-RING ubiquitin ligase complex genes in non-diseased and AD brains. In non-diseased brains, gene expression corresponding to protein targeting to the lysosome is reduced in this vulnerable population, while, in AD, we observed reduced expression of ribosome subunit genes. Genes linked to Aβ formation, including *BIN1*, *RTN4*, and *CASP3*, were upregulated in Exc-L2-3-*CUX2*-*CALB1* neurons in non-diseased brains, while those related to response to Aβ were upregulated in AD brains. We observed elevated expression of several tau-phosphorylating kinase pathways in both non-diseased and AD brains, such as those for extracellular signal-regulated kinase (ERK1/2) and calcium/calmodulin-dependent kinase (CAMK; Fig. 3a). We also found that Exc-L2-3-*CUX2*-*CALB1* neurons in non-diseased brains exhibit relatively higher expression of several tau-modifying kinase genes and lower expression of those for phosphatase subunits compared to other excitatory populations that we did not find to be significantly lost in AD (Fig. 3b). Gene expression differences notably also include upregulation of ERK1/2 family genes, *FYN*, and Rho-associated protein kinase (ROCK) genes and downregulation of genes encoding subunits of protein phosphatase 2A and 2B (PP2A, PP2B), and protein phosphatase 5 (PP5).

**Fig. 3.**
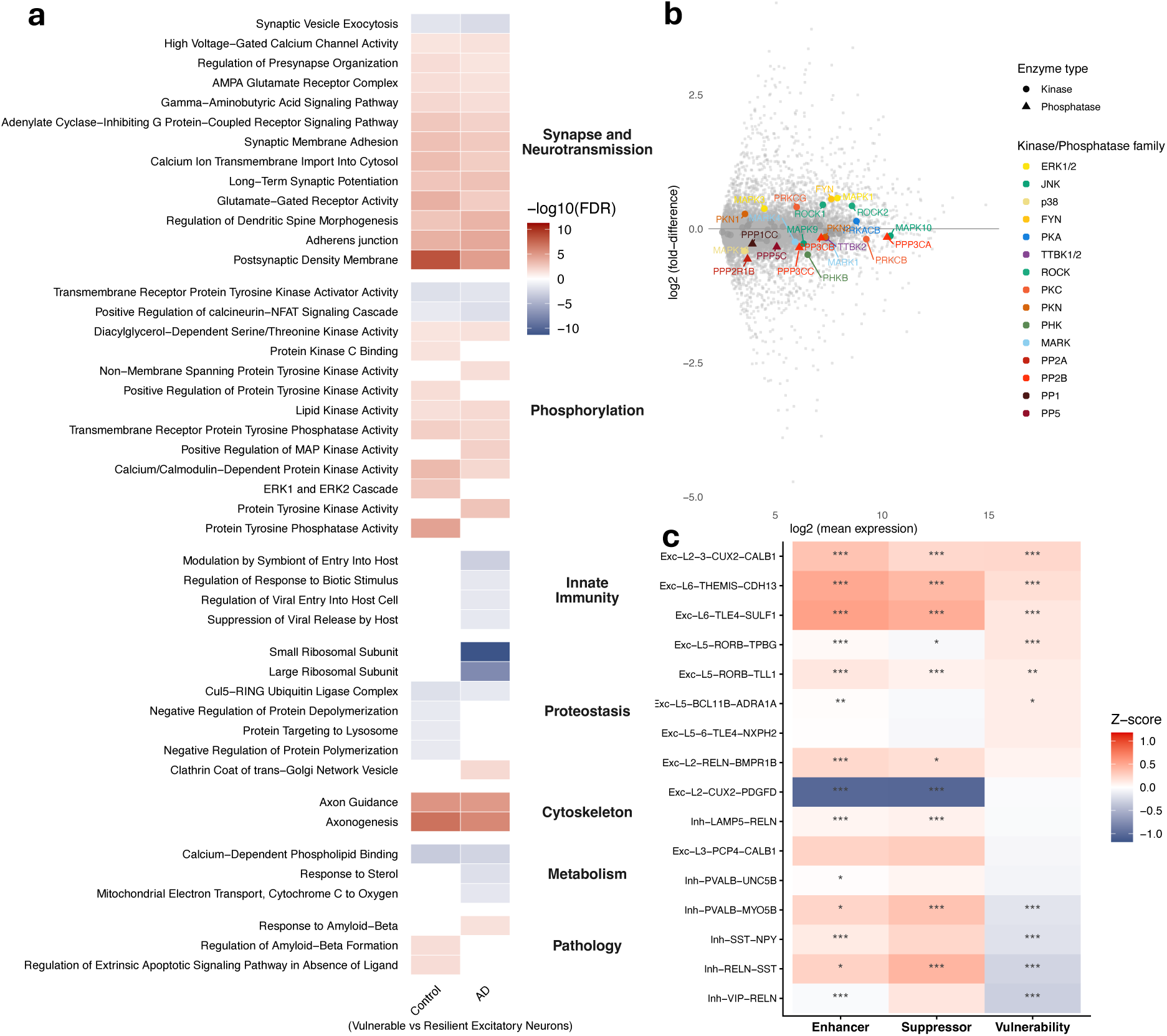
Intrinsic transcriptional determinants of neuronal vulnerability in entorhinal cortical neurons. a, Heatmap showing selected differentially expressed pathways in vulnerable versus resilient excitatory neurons of the EC, stratified by diagnosis (control and AD). n = 17 control donors, 19 AD; snRNA-seq cohort. **b,** MA plot (M, magnitude of change; A, average expression) of relative gene expression in vulnerable versus resilient excitatory neurons in non-diseased brains. Selected tau-associated kinases and phosphatases are highlighted by family. Points represent individual genes; only significantly differentially expressed genes (FDR < 0.05) are displayed. n = 17 control donors; snRNA-seq cohort. **c,** AUCell-derived scores for tau oligomerisation enhancer and suppressor gene sets^28^ and a composite vulnerability score (enhancer minus suppressor activity) across neuronal subclusters in non-diseased control donors. Higher vulnerability scores indicate greater inferred susceptibility to tau pathology. Heatmap displays z-scored module activity per subcluster. Subcluster differences were tested using one-versus-all linear mixed-effects models (score ∼ subcluster + age + sex + (1|donor)); p values were adjusted using the Benjamini-Hochberg method. *FDR < 0.05, **FDR < 0.01, ***FDR < 0.001. n = 17 control donors; snRNA-seq cohort.

We then tested for neuronal subtype differences in vulnerability to AD pathology using gene sets^28^ identified in CRISPR screens of iPSC-derived neurons to identify genes that enhance or suppress tau oligomerisation (Supplementary Table 11, 12). Using AUCell^29^, we scored transcriptomes of neurons from non-diseased brains for these gene sets and calculated a composite tau vulnerability score for each neuronal cluster. Exc-L2-3-*CUX2*-*CALB1* neurons exhibited the highest tau vulnerability score of all neuronal populations examined, while Inh-*VIP*-*RELN* neurons, amongst the most resilient to cell loss (Fig. 2a,c), exhibited the lowest (Fig. 3c; Supplementary Fig. 5; Supplementary Table 13). Rankings were largely preserved following the exclusion of samples expressing early AD-related pathology (Braak 2; Supplementary Fig. 6a,b).

### Intrinsic transcriptional vulnerability predicts tau pathology burden across neuronal subtypes

By comparing tau oligomerisation vulnerability scores of neurons from non-diseased brains with phospho-tau signal (pS396/pS404 sensitive PHF1 immunostaining) in matched IMC clusters (Supplementary Fig. 7a) from AD samples, we observed a moderate positive correlation (Fig. 4a; mean Spearman’s ρ = 0.388; p = 1.45 × 10^-10^), indicating that intrinsic vulnerability, as inferred from gene expression in non-diseased tissue, is reflected in downstream pathological change in AD. The correlation was robust to the exclusion of samples with early AD-related pathology (Braak 2) from the vulnerability scoring (mean Spearman’s ρ = 0.345; p = 3.97 × 10^-9^; Supplementary Fig. 6c). To characterise the gene-level basis of this association, we correlated relative expression of individual tau oligomerisation vulnerability genes across neuronal subtypes in non-diseased brains with the observed cluster-level PHF1 burdens in AD donors (Supplementary Fig. 7b; Supplementary Table 14). Genes were classified as concordant where the sign of the expression-PHF1 correlation matched the experimentally determined role of the gene in the CRISPR screen, and discordant otherwise. Gene set enrichment analysis (GSEA) uncovered significant enrichment of transcripts for respiratory electron transport pathways amongst the concordant tau oligomerisation suppressor genes expressed (Fig. 4b; Supplementary Table 15). A protein-protein interaction network analysis of the top 100 concordant suppressor genes revealed a dominant cluster of mitochondrial respiratory chain components spanning Complexes I (*NDUFB4, NDUFB7, NDUFB9, NDUFB11, NDUFS8, NDUFV2, NDUFAF6, NUBPL, TMEM126B, ACAD9*), III (*CYC1*) and IV (*COX5B, COX6B1, COX7B, COX7C, COX10, COX17, COA4*), mitochondrial ribosomal proteins (*MRPS9, MRPS14, MRPS31, MRPL19, MRPL33, MRPL39, GADD45GIP1*) and mitochondrial aminoacyl-tRNA synthetases (*AARS2, NARS2, WARS2, PARS2*; Fig. 4c). No equivalent pathway enrichment was identified for enhancer genes (Supplementary Fig. 7c). These findings implicate mitochondrial bioenergetic capacity as a determinant of neuronal resilience to tau phosphorylation.

**Fig. 4.**
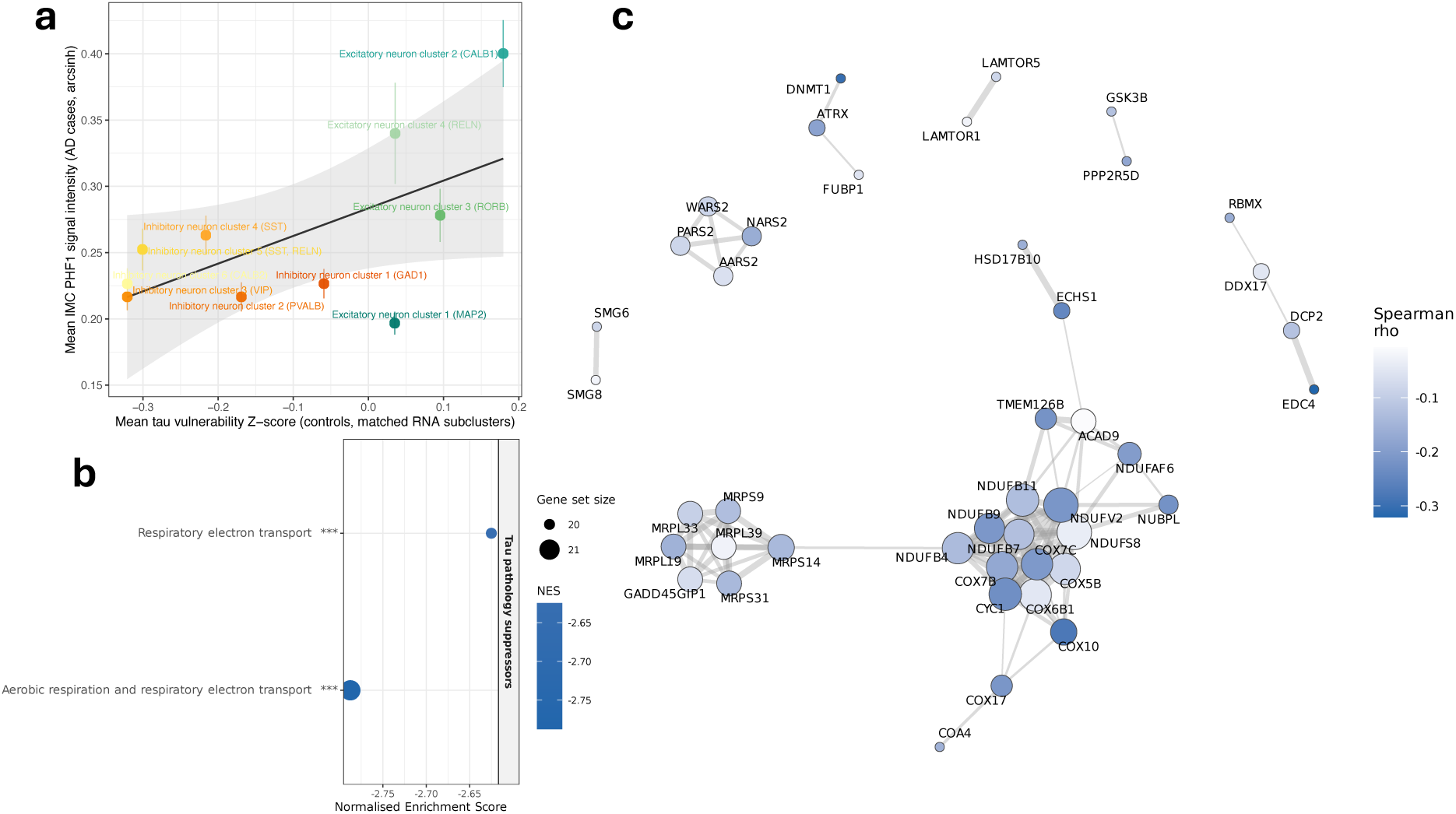
Validation and refinement of genetic predictors of tau pathology. a, Mean PHF1 signal intensity per IMC neuronal cluster plotted against tau vulnerability Z-score derived from non-diseased donor snRNA-seq. Each point represents a neuronal cluster (n = 10 clusters); error bars indicate SEM across AD donors. The regression line is fit to cluster means for visualisation only and does not reflect the inferential analysis (see text). Tau vulnerability scores and PHF1 measurements were obtained from independent cohorts (non-diseased controls and AD donors, respectively). Statistical inference was performed at the donor level: the Spearman rank correlation between cluster-level tau vulnerability Z-score and mean PHF1 was computed independently for each AD donor and per-donor correlations were Fisher z-transformed and tested against zero using a one-sample t-test (mean ρ = 0.388, 95% CI [0.301, 0.468]; t(43) = 8.37, p = 1.45×10⁻¹⁰; n = 44 AD donors). **b**, GSEA of concordant genes ranked by Spearman rho against the Reactome database. Only tau pathology suppressor genes reached significance; no enhancer pathways were significantly enriched. Dot size reflects gene set size; dot colour reflects normalised enrichment score (NES). Significance annotations reflect Benjamini-Hochberg-adjusted p-values (***p.adjust < 0.001). **c**, STRING protein-protein interaction network of the top 100 concordant tau pathology suppressor genes, ranked by magnitude of mean per-donor Spearman correlation with PHF1. Nodes are sized by network degree and coloured by Spearman rho.

### Calbindin-expressing excitatory neurons traverse a homeostasis-to-death trajectory

With IMC, we revealed that all neuronal clusters exhibited an increase in PHF1 signal with AD (adjusted p < 0.0001 for every cluster; Supplementary Table 16) but that excitatory cluster 2 (CALB1) and excitatory cluster 4 (RELN) exhibited steeper trajectories of tau phosphorylation than others that were associated with greater neuronal PHF1 burden in AD (Fig. 5a,b). To characterise neuron sub-type specific pathological burden in AD, we contrasted the abundance of 11 disease-associated protein markers, including those related to tau pathology, Aβ and impaired proteostasis in AD samples across neuronal subtypes (Fig. 5c). Phospho-tau labelled by AT8 (primarily targeting the pS202/pT205 epitope) and PHF1 (primarily targeting the pS396/pS404 epitope) was relatively elevated in excitatory cluster 2 (CALB1) neurons as well as in excitatory neuronal clusters 3 (RORB) and 4 (RELN). Excitatory cluster 2 (CALB1) neurons were distinct in also exhibiting elevated autophagosome protein LC3A/B immunostaining, consistent with either greater autophagic processing or impaired fusion and degradation of autophagosomes by lysosomes. Intracellular Aβ, as identified by MOAB-2, was elevated in excitatory cluster 4 (RELN), as well as in inhibitory clusters 2, 3, and 4. Increases in intracellular Aβ were not associated with significant neuronal loss in AD across these clusters.

**Fig. 5.**
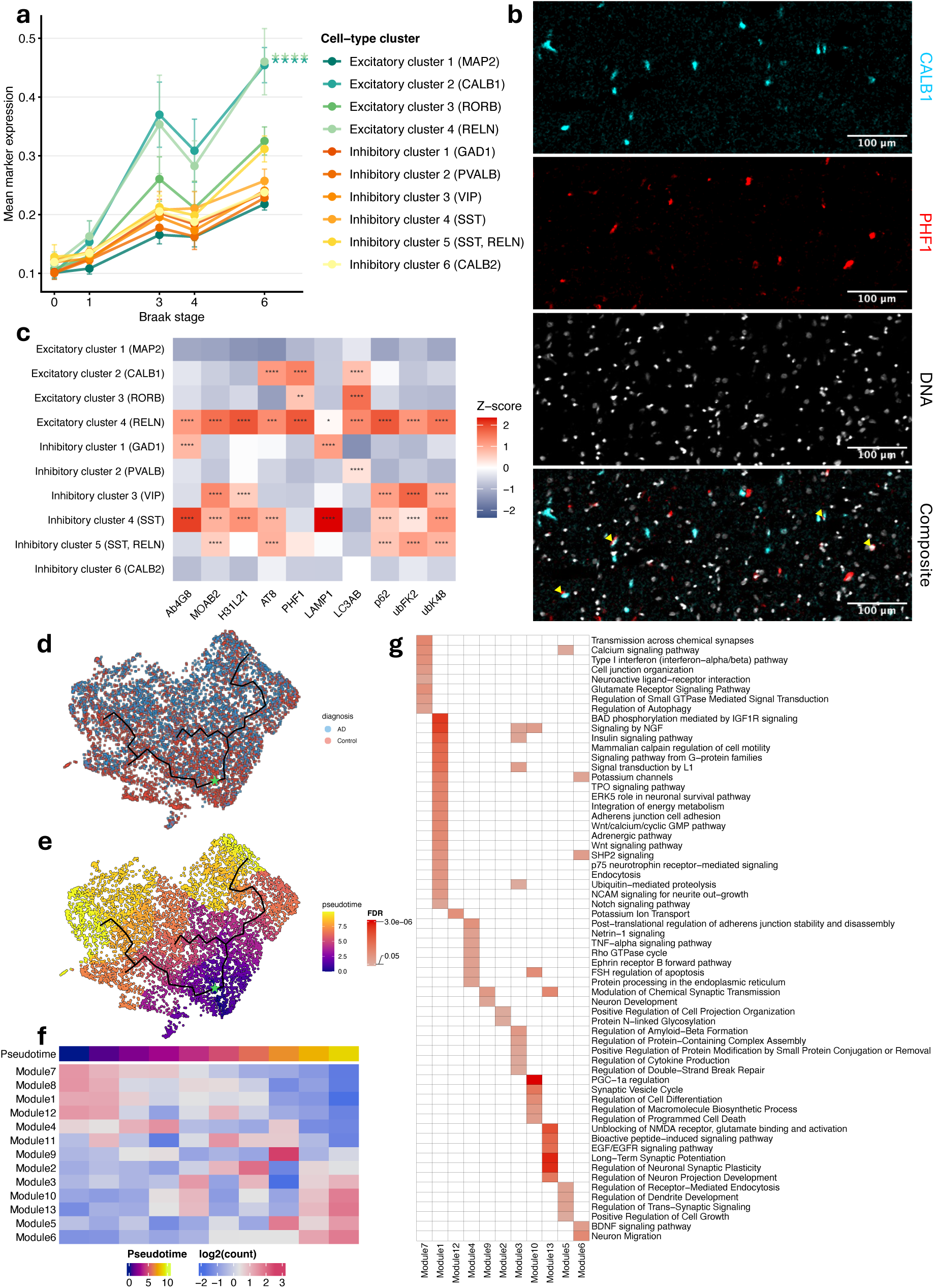
| Disease trajectory of vulnerable calbindin-expressing excitatory neurons. a, Mean PHF1 signal intensity across Braak stages for each IMC neuronal cluster. Points represent group means; error bars indicate SEM across donors. Differential disease trajectories were assessed per cluster (the “focal cluster”) based on linear mixed-effects models (mean PHF1 ∼ Braak × is_focal_cluster + age + (1 | donor)), where “is_focal_cluster” is a binary indicator (TRUE for the cluster being tested, FALSE for others). The “Braak × is_focal_cluster” interaction coefficient therefore estimates the difference in PHF1-versus-Braak slope between the focal cluster and the pooled remaining clusters, with a positive value indicating a steeper trajectory. P values were adjusted using the Benjamini-Hochberg method across clusters (****FDR < 0.0001). **b,** Representative IMC image from an AD donor, showing a region spanning layer 3 of the EC. PHF1 signal is shown in relation to calbindin-immunoreactive neurons, illustrating accumulation of phospho-tau within this population, with PHF1 signal also present in a subset of neighbouring calbindin-negative cells. **c,** Heatmap of AD-associated marker abundance across neuronal subtypes in AD samples. Statistical comparisons were performed using paired Wilcoxon tests (each cluster versus mean expression across all other neuron clusters); p values were adjusted using the Benjamini-Hochberg method (*p < 0.05, **p < 0.01, ***p < 0.001, ****p < 0.0001). **d,** UMAP of Exc-L2-3-CUX2-CALB1 neurons coloured by AD diagnosis. **e,** UMAP of Exc-L2-3-CUX2-CALB1 neurons coloured by pseudotime value; the trajectory root (green star) was set at the point of greatest proportional representation of non-diseased nuclei. **f,** Heatmap of gene expression module activity across pseudotime. **g,** Pathway enrichment of genes within each module, ordered by pseudotime progression. a: n = 18 control donors, 19 mid AD, 25 late AD; IMC cohort. c: n = 44 AD donors; IMC cohort. d-g: n = 17 control donors, 19 AD; snRNA-seq cohort.

Trajectory analysis of the vulnerable Exc-L2-3-*CUX2*-*CALB1* snRNA-seq neuronal population was performed to identify changes in transcriptional modules along the disease continuum. Nuclei were projected with UMAP and a trajectory was annotated, with the root being set as the node with the greatest proportion of neurons from non-diseased brains (Braak 0-1; Fig. 5d,e; Supplementary Fig. 8a-c). Gene expression was organised into discrete modules that peaked at different windows along the pseudotime trajectory (Fig. 5f). Modules with early pseudotime values (modules 7, 1, 12) were enriched for pathways involved in synaptic and trophic functions, including those related to synaptic transmission, calcium signalling, and neurotrophic pathways (NGF, IGF1R, Wnt; Fig. 5g; Supplementary Table 17). At intermediate pseudotimes (modules 4, 9, 2, 3), we observed a shift toward Aβ regulation, ubiquitin-mediated proteolysis, cytoskeletal regulation, and DNA double-strand break repair, suggesting an active cellular stress response in AD. Modules enriched at later pseudotimes (modules 10, 13, 5, 6) indicate loss of homeostasis with aberrant plasticity, developmental re-entry, and death priming (Fig. 5g). Overall, this trajectory analysis suggests that EC *CALB1*-expressing pyramidal neurons, which we find to accumulate phospho-tau and to be uniquely vulnerable to loss in AD, progress from a more homeostatic state characterised by expression of synaptic and trophic genes, through active responses to DNA damage and proteostatic stress, to a transcriptional signature consistent with neuronal dysfunction and early stages of cell death. The terminal modules thus provide a transcriptional correlate of the progressive neuronal loss that we observe by IMC.

### In-silico identification of modulators of EC neuronal resilience to tau pathology

Having defined a transcriptional axis distinguishing neurons resilient to tau pathology from those more vulnerable in the EC, we asked whether existing small molecules could be repurposed to push neurons toward the resilient end of this axis. We queried the LINCS L1000 compendium with the overlap of the concordant tau oligomerisation gene set and the genes that differentiate Exc-L2-3-*CUX2*-*CALB1* neurons from other excitatory neurons in non-diseased brains (Fig. 6a; Supplementary Table 18), restricting the query to results based on neural cell lines (neural progenitor cells (NPC), differentiated neurons (NEU), fibroblast-derived neural progenitor cells (FIBRNPC)) and filtering for blood-brain barrier (BBB) penetrance.

**Figure 6.**
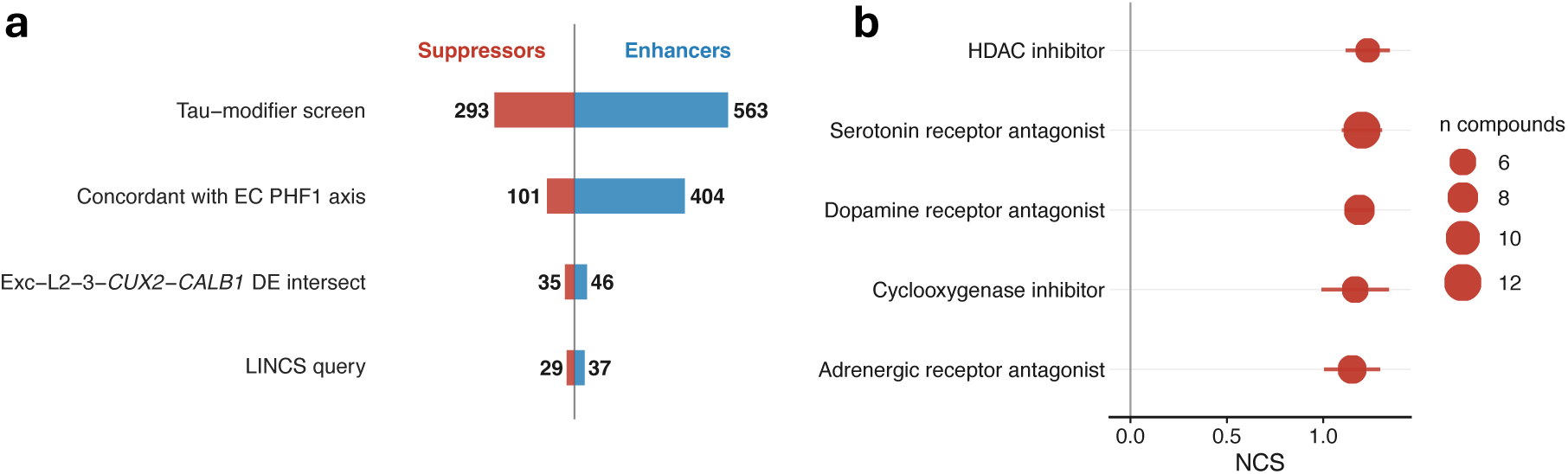
C**M**ap **analysis nominates mimickers of the EC neuronal tau-resilience axis. a,** Successive filters constructing the LINCS L1000 query (Suppressors, red; Enhancers, blue): tau-modifier screen (293/563); sign-concordance with per-donor IMC PHF1 burden across EC clusters (101/404); intersection with the Vulnerable-vs-Resilient DE signature of the Exc-L2-3-CUX2-CALB1 cluster (35/46); intersection with the LINCS phase I gene universe (final query: 29 up, 37 down; Supplementary Table 18). **b,** Top MoA classes by median compound NCS (top 5 mimickers). Compounds filtered for NCS > 1, FDR ≤ 0.05, and predicted BBB+ confirmed or likely; MoA classes restricted to those with 5 or more of such compounds. Dots mark per-class median NCS; horizontal segments span median ± median absolute deviation; dot area scales with the number of compounds in the class.

At the mechanism-of-action (MoA) level, several pharmacological classes showed coherent enrichment as mimickers of the resilience axis (Fig. 6b; Supplementary Fig. 9; Supplementary Table 19). The highest median normalised connectivity score (NCS) came from histone deacetylase (HDAC) inhibitors, followed by neuromodulator receptor antagonists (serotonin, dopamine, and adrenergic; occupying ranking 2, 3, and 5), and cyclooxygenase inhibitors, including meloxicam, the second highest-ranking pharmacologically annotated compound (NCS = 1.56, FDR = 2.16 × 10^-4^). Each class warrants validation in EC neuronal vulnerability models; the convergent signal across structurally unrelated chemotypes suggests the resilience program itself, rather than a specific target, is the drug-tractable axis.

## Discussion

The EC is amongst the earliest and most severely affected regions in AD^4,8^, yet the molecular identity of the most vulnerable neurons and mechanisms responsible for this vulnerability are still uncertain or poorly defined. Here, using convergent data from IMC and snRNA-seq, we have identified calbindin-expressing excitatory neurons as the most vulnerable population, demonstrating their preferential loss and phospho-tau accumulation with disease progression. The marker profile and laminar distribution of this population are consistent with the calbindin-expressing pyramidal neurons of EC layers 2 and 3 previously characterised by cytoarchitectonic and hodological studies in rodent and human tissue^24,30,31^. Transcriptomic profiling of these neurons in non-diseased brains reveals differences compared to more resilient neurons. This suggests an intrinsic susceptibility, characterised by relatively greater expression of several tau-modifying kinase pathway genes, lower expression of phosphatase pathway genes and an elevated tau oligomerisation vulnerability signature, is expressed in the layer 2 and 3 calbindin-expressing neurons even before the onset of AD-associated pathology. These findings thus focus attention on properties of neurons potentially modifiable through the life course as contributors to neuronal vulnerability in AD.

Early research established profound loss of superficial EC neurons in mild cognitive impairment^32^ and very mild AD^8^. Here, we identify a class of *CALB1*-expressing excitatory neurons in EC layers 2-3 as the most profoundly depleted neuronal population in our dataset. In rodents, these neurons project to hippocampal CA1^33^, and function as grid cells supporting spatial memory^34^ and temporal association memory^35^, with characteristic persistent firing patterns that impose chronic metabolic demands^36^. Grid-like spatial coding^37^ and high-precision temporal memory^35^ have both been demonstrated in the human EC, and selective loss of this population would be expected to disrupt both forms of memory processing early in AD^38,39^, consistent with the early clinical presentation. CA1 pyramidal neurons have independently been shown to be selectively vulnerable in AD^12,40^, raising the possibility that pathological vulnerability affects both the origins and targets of this direct memory circuit. Future work could explore whether quantitative measures of spatial and temporal memory could be sensitive biomarkers of dysfunction of these cells in early AD.

Our findings extend a recent large-scale single-cell transcriptomic study of AD^12^, providing molecularly resolved, spatially validated evidence for the specific excitatory subtype most susceptible to loss in the EC. The vulnerable Exc-L2-3-*CUX2*-*CALB1* population shows the highest transcriptomic similarity to the EC-specific cluster Exc L2/3 *TOX3 TTC6* reported previously^12^, which was significantly reduced in AD, suggesting that these clusters represent the same neuronal population. When the marker signature of Exc L2/3 *TOX3 TTC6* was transferred onto our excitatory neurons, the resulting population showed AD-associated reduction, validating our primary result.

Loss of calbindin-immunoreactive neurons in the EC has been reported in *post-mortem* AD studies since the early 1990s^9–11^, though these findings have not been widely integrated into current frameworks of EC vulnerability, in part due to concerns about the stability of the calbindin antibody epitope in stressed or dying neurons^41,42^. Our parallel IMC and snRNA-seq approach addresses this limitation by providing convergent evidence for calbindin neuron loss across modalities: snRNA-seq clustering was robust to *CALB1* depletion as the analysis was based on the expression of 594 marker genes with higher expression than *CALB1* within these neurons.

Two prior single-cell transcriptomic studies have reported proportional depletion of other EC neuronal subtypes, including *RORB*-expressing^13^ and *RELN*-expressing excitatory neurons^12^. Our data are consistent with pathological involvement of both, since excitatory clusters 3 (RORB) and 4 (RELN) showed relatively elevated phospho-tau accumulation by IMC, and excitatory cluster 4 (but not excitatory cluster 3) exhibited elevated intracellular Aβ; however, we found only trends towards cell loss of these neurons, without statistical significance, by either IMC or snRNA-seq.

How can the selective vulnerability of the Exc-L2-3-*CUX2*-*CALB1* population be explained? A key finding of our work is that *CALB1*-expressing EC neurons in non-diseased brains are distinguished from other excitatory neurons by their greater relative expression of tau-modifying kinase pathways and lower phosphatase expression. This may prime them for pathological tau phosphorylation. We identified evidence that serotonin, dopamine, and adrenergic receptor antagonists mimicked the resilience module for calbindin neurons. These neuromodulators stimulate cAMP-mediated PKA activation, which supports tau phosphorylation^43^. At the gene level, overexpressed kinases include ERK1/2, FYN, and ROCK, while downregulated phosphatases include PP2A, PP2B, and PP5 – all with well-established roles in tau modification. Each kinase carries distinct disease relevance: FYN phosphorylates tau at tyrosine 18 and mediates tau-dependent excitotoxicity downstream of Aβ42^44^; ERK1/2 activation has been reported in neurons bearing neurofibrillary tangles in the human AD brain^45^; and ROCK promotes tau aggregation and amyloid processing, and has emerged as a candidate therapeutic^46^. PP2A loss of function is a consistently replicated molecular finding in AD brain tissue^43,47^. We propose that the co-occurrence of these features in non-diseased tissue represents a compounded, “two-hit” susceptibility.

Beyond the kinase/phosphatase imbalance, our CRISPR screen-derived vulnerability scoring revealed additional susceptibility factors, with the vulnerable Exc-L2-3-*CUX2*-*CALB1* cluster scoring highest for tau oligomerisation vulnerability of all EC populations examined. Gene-level decomposition revealed that neuronal resilience to tau phosphorylation is associated with the level of expression of mitochondrial respiratory chain component genes, consistent with prior work showing that mitochondrial dysfunction promotes tau hyperphosphorylation through impaired PP2A activity^47,48^ and metabolic stress-mediated kinase activation^49,50^. Lower mitochondrial bioenergetic capacity may therefore contribute to the intrinsic susceptibility of *CALB1*-expressing EC neurons to neurodegeneration in AD.

Differences in expression of proteostatic and synaptic gene sets further distinguished these neurons. Relatively reduced expression of genes encoding the Cul5-RING ubiquitin ligase complex, lysosomal protein targeting machinery, and ribosomal subunits suggests a lower capacity for upregulating protein turnover, a hypothesis supported by the observation by IMC of greater LC3A/B in calbindin-positive neurons with AD. Reduced expression of presynaptic vesicle exocytosis genes alongside a relative enrichment of postsynaptic glutamate receptor and long-term potentiation pathways, and calcium channel expression provides additional susceptibility factors in the context of disease-related triggers to excitotoxicity^51^.

The vulnerability of calbindin-expressing neurons may appear paradoxical given the neuroprotective role of calbindin as an intracellular calcium buffer^52^ and reports that calcium-binding-protein-expressing interneurons are relatively spared in AD^53,54^. However, the calcium buffering capacity of calbindin may be overwhelmed by the sustained calcium load imposed by the abnormal, increased activity of these neurons, particularly if expression is downregulated in stressed neurons with disease^41,42^. Moreover, calbindin expression is just one characteristic of these neurons; we have identified others that may promote neurodegeneration.

Trajectory analysis revealed a transcriptional progression towards neurodegeneration with AD in the calbindin neurons. At early pseudotimes, modules enriched for synaptic transmission, calcium signalling, and neurotrophic survival (NGF, IGF1R, Wnt) defined a homeostatic state; progressive loss of these signatures was associated with transcriptional markers of early synaptic dysfunction associated with neuronal loss in AD^55,56^. At intermediate pseudotimes, a shift toward increased expression of ubiquitin-mediated proteolysis, cytoskeletal dysregulation, Aβ regulation, and DNA double-strand break repair pathways suggests an active cellular stress response. Impaired DNA damage repair may drive neurodegeneration in AD^57^. At later pseudotimes, co-expression of modules enriched for aberrant developmental plasticity, neuronal projection development, BDNF signalling, and programmed cell death pathways reflects re-activation of developmental programmes and suggests failure of adaptive responses or pathological cell cycle re-entry, which has been associated with neuronal death rather than regeneration^58–60^. The co-occurrence of these signatures suggests that late-stage vulnerable neurons are transcriptionally active but functionally compromised, supporting a prolonged degenerative state rather than an abrupt transition.

Halting this pathological cascade, or preventing its initiation, requires shifting vulnerable neurons towards more resilient transcriptional states. We queried the LINCS L1000 dataset with the resilience signature to nominate small molecules predicted to engage this axis. HDAC inhibitors emerged as the drug class most strongly aligned with the resilience signature, consistent with a previous *in silico* drug screen^61^, and the reported elevation of HDAC2 in AD brain tissue and in mouse models of neurodegeneration^62^. In a mouse model of AD, HDAC inhibition was shown to rescue memory deficits^63^. Cyclooxygenase inhibitors, including meloxicam, also reproduced the resilience signature, corroborating epidemiological evidence linking long-term NSAID use to reduced AD incidence^64,65^. The ADAPT trial found no clinical benefit in symptomatic AD^66^, but there was evidence for prevention of AD in pre-symptomatic individuals^67^. The preferential COX-2 selectivity of meloxicam is notable because COX-2 is the activity-inducible prostaglandin synthase constitutively expressed in cortical and hippocampal neurons and upregulated in AD brains^68,69^, and a known modulator of glutamatergic excitability and intracellular calcium handling^70^. Three of the top five suggested therapeutic classes by mechanism of action were serotonin, dopamine, and adrenergic signalling antagonists. That three structurally and mechanistically distinct receptor classes converged onto the same transcriptional output suggests the relevant pharmacological lever is not a specific receptor but a shared downstream response that secondarily engages mitochondrial biogenesis and proteostatic capacity.

Beyond insights from the LINCS L1000 dataset, resilience mechanisms also could be enhanced by saracatinib (AZD0530), a FYN kinase inhibitor that showed a trend to slowing EC degeneration by MRI in an early phase clinical trial^71^, or sodium selenate, a PP2A activator that also has proven safe in small, early clinical studies^72^. Fasudil, a ROCK1/2 inhibitor, reduced phospho-tau in a mouse model^73^ and an early-stage clinical trial is underway (NCT06362707). Our results also support the investigation of mitophagy activators such as urolithin A, currently a focus of clinical investigations for aging and metabolic disease^74^.

Limitations of our work should be acknowledged. First, these data were derived from *post-mortem* human brain tissue, which precludes causal inference, and we cannot fully exclude contributions of age-related or other pathological processes to the transcriptional signatures we have defined. Second, not all IMC neuronal nuclei or soma were assigned to specific subtypes. The ratio of excitatory-to-inhibitory neurons was lower than previously estimated^75^, highlighting that some excitatory populations may have been underrepresented in the analysis, although the unclassified nuclei or cells did not appear to be amongst those with a trend to loss in AD. Third, the pseudotime trajectory analysis provides an inferred transcriptional continuum across different samples rather than a longitudinal measure of disease. A specific factor that could confound interpretation of this would be differential enrichment of samples for *APOE4* across Braak stages. *APOE4* alleles are independently associated with impaired ubiquitin-mediated proteolysis, cytoskeletal dysregulation, Aβ regulation, and DNA double-strand break repair pathways^76–79^. Based on genotyping for 27 of 36 snRNA-seq samples, *APOE4* allele carriers were 2.1-fold more common in Braak 5-6 relative to Braak 0-2 donors. Future work needs to test whether the calbindin neurons are uniquely sensitive to these effects of *APOE4*. Fourth, while identification of the snRNA-seq calbindin cluster could be done independently of *CALB1* expression, identification of the corresponding IMC cluster could not. This is in part due to the limited number of excitatory neuron subtype markers in the panel, which restricted excitatory neuron annotation with IMC. The joint evidence for calbindin neuron loss, although complementary, is confined by potential ambiguity concerning effects of any reduced *CALB1* expression rather than neuronal loss. Finally, both the iPSC-derived neuron CRISPR screen and the LINCS L1000 neural reference profiles derive from developmentally immature, regionally unspecific cells, and the resulting inferences should be treated as hypothesis-generating pending validation in aged EC-relevant models.

In summary, we identified calbindin-expressing excitatory pyramidal neurons of EC layers 2-3 as the primary vulnerable population in AD using convergent data from IMC and snRNA-seq. We found evidence that factors contributing to their susceptibility are rooted in intrinsic molecular features present before the onset of disease. These include a kinase/phosphatase imbalance favouring tau hyperphosphorylation and a lower mitochondrial bioenergetic capacity. The resilience axis defined by these features provides a substrate for drug repurposing or new therapeutics to promote more resilient transcriptional states based on evidence from studies of HDAC inhibitors, cyclooxygenase inhibitors, and antagonists of several neuromodulator receptors to date. The suggestion of mechanistic synergy related to the “two-hit” pathology cascade we identified specifically supports the consideration of combination therapies.

## Methods

### Tissue samples

This study was carried out in accordance with the Regional Ethics Committee and Imperial College London Use of Human Tissue guidelines. Samples were selected based first on neuropathological diagnosis (non-AD or AD) from four UK brain banks (Oxford Brain Bank [University of Oxford], South West Dementia Brain Bank [University of Bristol], Cambridge Brain Bank [University of Cambridge], and Multiple Sclerosis and Parkinson’s Tissue Bank [Imperial College London]) and the Netherlands Brain Bank. We excluded cases with clinical or *post-mortem* diagnosis of potentially confounding neurological conditions, including but not limited to Parkinson’s disease, Huntington’s disease, progressive supranuclear palsy, epilepsy, hippocampal sclerosis, frontotemporal dementia, age-related tau astrogliopathy, argyrophilic grain disease, and traumatic brain injury. Further exclusion criteria included type 2 diabetes mellitus, the presence of infarcts greater than 10mm in diameter unless confined to the cerebellum, cerebrovascular disease (such as cerebral atherosclerosis, small vessel disease, arteriosclerosis) described as moderately severe to severe, and a *post-mortem* interval greater than 41 hours. IMC samples comprising the non-AD group were all required to have no report of dementia or cognitive impairment, *post-mortem* confirmation of the absence of AD pathology, and a Braak stage of 0 or 1. Samples within the mid-stage AD group all had a clinical diagnosis of dementia before death and *post-mortem* confirmation of AD pathology defined as Braak stages 3 or 4. The late-stage group only included Braak stage 6 samples with a clinical dementia diagnosis before death. The final IMC cohort was formed of human post-mortem FFPE entorhinal cortex slices of 8μm thickness from 23 non-diseased controls, 21 individuals with mid-stage AD, and 26 with late-stage AD (Fig. 1a). For single nucleus RNA sequencing, frozen entorhinal cortex tissue was available for 19 matched samples (9 non-AD, 4 mid-stage AD, 6 late-stage AD). These frozen blocks were supplemented with 2 additional mid-stage AD cases to increase statistical power and combined with a previously described cohort of 19 entorhinal cortex snRNA-seq samples^22^. All the brain banks used have generic research ethics committee approval to function as research tissue banks and therefore did not require additional ethics panel approvals for use by UK researchers.

### Designing and testing of the neuronal antibody panel for imaging mass cytometry (IMC)

A list of candidate markers for excitatory and inhibitory neurons, microglia, astrocytes, and AD-associated proteins was formed based on a previous study by our group^20^ and supplemented with additional markers for neurons enriched in the EC^21^ and previously identified vulnerable neurons^13,18^.

All antibodies in the final panel were validated using immunofluorescence to confirm expected subcellular localisation and expression patterns and subsequently optimised for IMC. The final antibody panel (Table 1) included two pan-neuronal markers (HuC/D, MAP2), 16 neuronal subtype-specific markers (CUX2, Calretinin [CALB2], LMO4, Calbindin [CALB1], PCP4, VIP, GPC5, FOXP2, PVALB, RORβ, SST, NPY, GAD1, Reelin [RELN], ADARB1, LHX6), 5 glial markers (Iba1, CD68, S100b, GFAP, OLIG2), 3 Aβ markers (4G8, MOAB-2, H31L21), 2 phospho-tau markers (PHF1, AT8), and 6 markers of proteostasis (ubK48, ubFK2, LC3A/B, p62, LAMP1, BNIP3). Nuclei were counterstained with a natural-abundance iridium nucleic acid intercalator (Cell-ID Intercalator-Ir, Standard BioTools). CCK was included in the original panel design but excluded from the study due to supply issues. OLIG2 was not included in the analysis due to unreliable signal.

**Table 1.** Final panel of IMC antibodies.

| Antibody | Gene | Supplier | Catalogue | Conjugate | Dilution |
| --- | --- | --- | --- | --- | --- |
| **ub K48** |  | Abcam | ab271911 | 141 Pr | 1:200 |
| **CUX2** | *CUX2* | Abnova | H00023316-M03 | 143 Nd | 1:50 |
| **Aβ (4G8)** | *APP* | BioLegend | 800702 | 144 Nd | 1:1000 |
| **Calretinin** | *CALB2* | Abcam | ab232462 | 145 Nd | 1:500 |
| **pTau (PHF1)** | *MAPT* | Peter Davies Laboratory, Albert Einstein College of Medicine |  | 146 Nd | 1:500 |
| **LC3AB** | *MAP1LC3A/B* | Cell Signalling | 58139 | 147 Sm | 1:200 |
| **HuCD** | *ELAVL3/4* | Thermo Fisher Scientific | A21271 | 148 Nd | 1:200 |
| **LMO4** | *LMO4* | Thermo Fisher Scientific | PA5-19062 | 149 Sm | 1:50 |
| **ub FK2** |  | Merck | 04-263 | 150 Nd | 1:200 |
| **LAMP1** | *LAMP1* | Fluidigm | 3151021D | 151 Eu | 1:200 |
| **CALB1** | *CALB1* | Abcam | ab233018 | 152 Sm | 1:300 |
| **Aβ (H31L21)** | *APP* | Thermo Fisher Scientific | 700254 | 153 Eu | 1:200 |
| **Aβ (MOAB2)** | *APP* | Biosensis | M-1586-100 | 154 Sm | 1:100 |
| **PCP4** | *PCP4* | Sino-biological | 207078-T08 | 155 Gd | 1:50 |
| **OLIG2** | *OLIG2* | Abcam | ab220796 | 156 Gd | 1:120 |
| **VIP** | *VIP* | Abcam | ab273589 | 158 Gd | 1:50 |
| **GPC5** | *GPC5* | LSBio | LS-C166601-400 | 159 Tb | 1:50 |
| **MAP2** | *MAP2* | Abcam | ab236033 | 160 Gd | 1:1500 |
| **FOXP2** | *FOXP2* | Sino-biological | 204066-T08 | 161 Dy | 1:50 |
| **GFAP** | *GFAP* | Abcam | ab218309 | 162 Dy | 1:500 |
| **PVALB** | *PVALB* | NovusBio | NB120-11427 | 163 Dy | 1:1000 |
| **p62** | *SQSTM1* | Abcam | ab227992 | 164 Dy | 1:200 |
| **pTau (AT8)** | *MAPT* | Thermo Fisher Scientific | MN1020 | 165 Ho | 1:200 |
| **RORB** | *RORB* | Sino-biological | 202727-T08 | 166 Er | 1:200 |
| **S100B** | *S100B* | NovusBio | NBP2-54426 | 167 Er | 1:1000 |
| **SST** | *SST* | Abcam | ab108456 | 168 Er | 1:200 |
| **Iba1** | *AIF1* | WAKO | 019-197471 | 169 Tm | 1:3000 |
| **NPY** | *NPY* | LSBio | LS-B6400 | 170 Er | 1:650 |
| **BNIP3** | *BNIP3* | Novus Biological | NBP1-77683 | 171 Yb | 1:50 |
| **GAD1** | *GAD1* | Abcam | ab240280 | 172 Yb | 1:200 |
| **RELN** | *RELN* | Santa Cruz | sc-25346 | 173 Yb | 1:50 |
| **CD68** | *CD68* | Abcam | ab227458 | 174 Yb | 1:60 |
| **ADARB1** | *ADARB1* | LSBio | LS-C176789 | 175 Lu | 1:55 |
| **LHX6** | *LHX6* | LSBio | LS-B10690 | 176 Yb | 1:55 |
| **nuclei** |  | Fluidigm | 20192B | 191 Ir | 1:400 |

### Immunofluorescence staining and confocal acquisition

8μm sections were cut from an FFPE optimisation block using a microtome and placed over SuperFrost Plus slides for antibody optimisation and validation. All slides were baked at 60°C for at least one hour before staining to allow FFPE sections to adhere to the slide. Sections were deparaffinised with two consecutive 5-minute incubations with xylene, rehydrated with a graded ethanol series (100%, 100%, 90%, 70%) for 5 minutes each, and washed in distilled water (dH_2_O) for 5 minutes. Antigen retrieval was performed by incubating slides in pH 9.0 ethylenediaminetetraacetic acid (EDTA) for 20 minutes in a steam chamber, then slides were cooled on ice for 10 minutes and washed in dH_2_O then in phosphate-buffered saline (PBS) for 5 minutes each. A hydrophobic barrier was drawn around the tissue using a PAP pen (Abcam) immediately before slides were incubated in 200μl blocking buffer consisting of 10% donkey serum (Abcam) in PBS with 0.3% Triton X-100 for 1 hour at room temperature inside a humid chamber. Primary antibodies were diluted in blocking buffer and applied to slides overnight in a humid chamber at 4°C. Slides were then washed in PBS for 5 minutes 3 times before applying Alexa Fluor cross-absorbed secondary antibodies (Invitrogen) and 4’,6-diamidino-2-phenylindole (DAPI) diluted 1:500 in PBS and incubated for 1 hour at room temperature in a dark humid chamber. Slides were then washed twice with dH_2_O for 5 minutes each, incubated for 1 minute with TrueBlack autofluorescence quencher (Biotium) diluted to 1× in 70% ethanol, then washed 3 further times in dH_2_O for 5 minutes each. Glass coverslips were mounted with ProLong^™^ Diamond Antifade Mountant (Invitrogen). Stained sections were acquired with a Leica TCS SP8 confocal microscope at 20× magnification.

### Antibody conjugation, staining, and acquisition of IMC

Primary antibodies were required to be free of carriers such as bovine serum albumin (BSA), which can interfere with the conjugation protocol. Primary antibodies that were not suspended in carrier-free solution (LMO4 and RELN) were purified using antibody purification kits (Protein A, Abcam) following manufacturer instructions. Carrier-free and purified antibodies were then conjugated to lanthanide metals using the MaxPar V8 kit (Fluidigm, Standard BioTools) according to the manufacturer instructions.

After optimisation and validation of the conjugated antibodies, one FFPE section per sample was stained with the full conjugated antibody panel (Table 1). Samples underwent deparaffinisation, rehydration, antigen retrieval, and blocking as in the immunofluorescence protocol described above. After blocking, the entire conjugated antibody panel was diluted in blocking buffer alongside the nucleic acid intercalator and 80μl of the diluted panel solution was applied to each slide. Tissue area was adjusted to a maximum of 2cm height and 3cm width to maintain consistent antibody coverage across all samples. Slides were incubated overnight in a humid chamber at 4°C. Slides were washed in dH_2_O for 10 minutes 3 times the following day and air dried for at least 20 minutes. IMC was performed using a Hyperion Tissue Imager coupled to a Helios mass cytometer (Fluidigm, Standard BioTools). The instrument was tuned using the manufacturer’s 3-Element Full Coverage Tuning Slide before the slides were loaded into the device. 3 ROIs from medial to lateral EC were ablated from the same FFPE section for each sample at a laser frequency of 200Hz with 1μm resolution. Each ROI spanned the full cortical depth from the pial surface to the white matter boundary, totalling 6 hours of acquisition per sample (∼2 hours per ROI, corresponding to 1.21 ± 0.04 mm^2^ of area ablated per ROI). Data were saved as .txt files for subsequent processing.

### IMC nuclei segmentation and pre-processing

Steinbock^80^ (v0.16.3) was used for image processing and quantification of protein signal intensity in segmented nuclei. A panel csv was extracted from .txt files and was formatted manually to select which channels should be used. Image files (.tiff) were produced and a hot pixel filter (threshold = 50) was applied. Images were extracted and DNA masks were created using a FIJI ImageJ^81^ (v2.14.0) macro. The Phansalkar local thresholding algorithm (radius = 50μm) was employed to distinguish true nuclear signal from the background. After thresholding, a watershed transformation separated touching nuclei, and a minimum nucleus area of 16μm^2^ was set. With the Steinbock measure intensities command, signal intensities for all markers were measured within each segmented nucleus. Nuclei morphology features were exported, and cellular neighbours were recorded as those overlapping following a 10μm expansion.

Downstream IMC analysis was performed in R (v4.4.2; Supplementary Table 20) using published scripts^80^ and custom additions. Marker intensities were transformed using the arcsinh function (cofactor = 1) and signal spillover between channels was quantified and mathematically corrected for using the CATALYST R package^82^.

### IMC quality control

245,497 nuclei across 209 ROIs from 70 donors were taken into quality assessment. Fourteen ROIs were excluded through unbiased visual assessment of images. After this, no ROIs met the exclusion criteria of having nuclear density outside of 2.5 standard deviations from the interquartile boundaries. 15,082 nuclei were removed from the analysis due to having no recognisable marker signal. This was determined by fitting a Gaussian mixture model for every marker within every sample to identify background and foreground signal. Nuclei were excluded if they had no signal above the background threshold across all markers. For statistical comparison of cell types with AD pathology, we only included samples with all 3 ROIs remaining (18 non-diseased controls, 19 mid-stage AD, 25 late-stage AD).

### IMC cell phenotyping

Cell phenotyping was performed in two stages. First, PhenoGraph clustering^83^ (Rphenograph; k = 50) was applied to the Harmony-corrected low-dimensional embedding of all cells to resolve major cell lineages. Community detection was performed using the Louvain algorithm. Cluster annotations were assigned based on differential marker abundance identified using Seurat’s FindAllMarkers^84^ function (Wilcoxon rank-sum test; minimum cell fraction = 0.2; log2FC threshold = 0.2) and validated by pseudobulked mean expression profiles per cluster. Twenty distinct clusters were identified; one cluster (Artefact cluster) was identified as an artefact based on non-specific co-expression of multiple lineage markers and excluded from all downstream analyses. Broad cell type classes were assigned based on canonical marker profiles (microglia: IBA1+, CD68+; astrocytes: GFAP+, S100β+; excitatory neurons: MAP2+, CALB1+, RORβ+; inhibitory neurons: GAD1+, PVALB+, VIP+, SST+).

In the second stage, cells assigned to neuronal lineages from stage one were isolated and re-embedded independently. PCA was performed on 14 neuronal subtype markers (RORβ, GAD1, RELN, PCP4, CUX2, VIP, SST, NPY, PVALB, CALB2, FOXP2, LMO4, LHX6, CALB1), retaining 8 principal components. Harmony correction was re-applied to the neuronal PCA space (grouping by donor), and UMAP embedding was computed using the Harmony-corrected components (n_neighbours = 7, min_dist = 0.001, spread = 1.5, metric = Euclidean). PhenoGraph clustering was then applied to the neuronal Harmony embedding (k = 45), yielding 17 neuronal subclusters. Subclusters were annotated using the same FindAllMarkers approach as stage one, and biologically related subclusters with overlapping marker profiles were merged, yielding 4 excitatory and 6 inhibitory neuronal populations. Neuronal subcluster annotations were then projected back onto the full-dataset UMAP for visualisation.

### Analysis of columnar cell densities

Columnar cell densities were calculated as the number of nuclei per mm along the pial surface of the cortex and spanning the depth of the grey matter to the white matter boundary. The columnar density is explicitly the thickness-adjusted areal density used to account for variation in cortical thickness. This method allows consistent quantification between samples^25^, irrespective of brain curvature^26^.

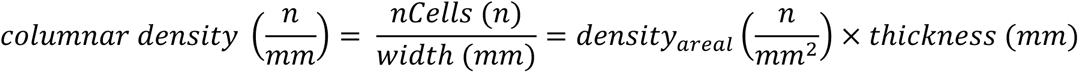

To assess disease-associated changes in columnar density, a linear mixed effects model was fitted independently for each cell type cluster using the lmerTest R package^85^, with the model formula: columnar density ∼ AD status + Age + (1|patient_id), where AD status was defined based on clinical diagnosis, and patient_id was included as a random effect to account for the non-independence of multiple ROIs from the same donor. Only donors with all three ROIs passing QC were included. *APOE* genotype was not applied as a covariate due to the genotype being unknown for 33 of the 62 samples analysed. P values were adjusted for multiple comparisons across cell type clusters using the Benjamini-Hochberg method. For visualisation, the mean percentage change in columnar density relative to the control group mean was calculated per cluster across AD donors.

To characterise columnar density across disease severity in clusters of interest, a separate linear mixed-effects model was fitted for each cluster using Braak stage group as a three-level factor (Braak 0-1, Braak 3-4, Braak 6): columnar density ∼ BraakGroup + Age + (1|patient_id). P values were adjusted using the Benjamini-Hochberg method within broad cell type groups.

### Characterisation of cell-type-specific marker trajectories

To characterise cell-type-specific trajectories of PHF1 (phospho-tau S396/S404) signal intensity across disease progression, PHF1 intensity values from the arcsinh-transformed IMC expression matrix were aggregated to ROI-level means per neuronal cluster, retaining only donors with all three ROIs passing quality control. To identify clusters with differential disease trajectories relative to all other neuronal clusters combined, a linear mixed-effects model was fitted independently for each neuronal cluster using the lmerTest R package, with the model formula: mean PHF1 ∼ Braak × is_focal_cluster + Age + (1|patient_id), where Braak stage was treated as a continuous predictor, is_focal_cluster is a binary indicator for the cluster under test, and patient_id is included as a random intercept to account for the non-independence of multiple ROIs from the same donor. The interaction coefficient (Braak:is_focal_clusterTRUE) quantifies the degree to which a given cluster’s PHF1 trajectory across Braak stages is steeper or shallower than that of all remaining neuronal clusters combined. Clusters with a significantly positive interaction coefficient were identified as exhibiting disproportionately steeper phospho-tau accumulation with disease progression. P values were adjusted for multiple comparisons across clusters using the Benjamini-Hochberg method.

### Characterisation of cell-type-specific marker abundance

To characterise the abundance of AD-associated pathological markers across neuronal subtypes, arcsinh-transformed IMC marker intensities for state markers (Aβ: 4G8, MOAB-2, H31L21; phospho-tau: AT8, PHF1; proteostasis: LAMP1, LC3A/B, p62, ubFK2, ubK48) were extracted for all neuronal cells from AD donors, excluding donors with fewer than three ROIs passing quality control. Marker intensities were averaged first across cells within each ROI per cluster, then across ROIs within each donor, to produce donor-level mean intensities per marker per cluster. To identify neuronal clusters with selectively elevated marker abundance, a paired one-sided Wilcoxon test was performed for each marker-cluster combination, comparing the donor-level mean intensity within the cluster under test against the mean intensity within all other neuronal clusters combined, calculated as the leave-one-out mean across clusters per donor. P values were adjusted for multiple comparisons across all marker-cluster combinations using the Benjamini-Hochberg method. For visualisation, cluster-level mean expressions were z-scored within each marker across clusters, producing a z-score matrix displayed as a heatmap with significance annotations overlaid.

### Isolation of nuclei for single nucleus RNA sequencing

Single nucleus isolation was conducted on 19 samples from the IMC cohort (9 controls, 4 mid-stage AD, 6 late-stage AD) plus 2 additional mid-stage AD samples to enhance statistical power. Fresh frozen human entorhinal cortex tissue blocks were cryosectioned at 80μm and 50mg of tissue was collected in an RNase-free Eppendorf tube. Nuclei were isolated based on published protocols^86,87^. All steps were carried out on ice or at 4°C. Tissue was homogenised in 1.2ml homogenisation buffer (250mM sucrose + 25mM KCl + 5mM MgCl_2_ + 10mM pH9 Tris Buffer + 1mM DTT + 1× protease inhibitor cocktail (Promega) + 1U/μl RNasin RNase inhibitor (Promega) + 0.1% Triton X-100 + 0.1% DAPI) using a 2ml glass dounce. The homogenate was centrifuged for 8 minutes at 500g, and the supernatant was removed. The homogenate was resuspended in homogenisation buffer and filtered through a 70μm filter followed by an iodixanol density gradient (29%, 50%) centrifugation at 13,000g for 40 minutes. The pellet was resuspended in 500μl BSA buffer (1% BSA, 1U/μl RNasin RNase inhibitor, 1× protease inhibitor cocktail, PBS), filtered through a 35μm nylon mesh and centrifuged at 500g for 5 minutes. The nuclei were then resuspended in 1ml BSA buffer and filtered again into a 5 ml FACS tube. 10μl of suspended nuclei was loaded onto a cell imaging slide and taken for visual inspection at 20× on a microscope. Nuclei were sorted using a FACS Melody (BD Bioscience) to filter for DAPI-positive single nuclei.

### Single nucleus processing and sequencing

A 9μl aliquot of isolated nuclei was stained with 1μl propidium iodide and counted on a LUNA-FL Dual Fluorescence Cell Counter (Logos Biosystems). Sufficient nuclei for recovery of 9000 cells were used for 10x Genomics Chromium Single Cell 3’ processing and library generation. All steps were conducted according to the 10x Genomics Chromium Single Cell 3’ Reagents Kits v3.1 User Guide, with the exception that 10 cycles of cDNA amplification were performed and 25ng of amplified cDNA per sample was taken through for fragmentation. The final index PCR was conducted at 14 cycles. cDNA and library preparation concentration were measured using the Qubit dsDNA High Sensitivity Assay Kit (Thermo Fisher Scientific), and library preparations were assessed using the TapeStation D5000 ScreenTape Assay (Agilent). Pooled samples at equimolar concentrations were sequenced on a NovaSeq6000 according to the standard 10x Genomics protocol.

### Pre-processing and quality control of snRNA sequencing data

Sequencing reads were demultiplexed and aligned to the GRCh38 reference. Cellbender remove-background v3^88^ with the default settings was applied to reduce background noise from ambient RNA and index hopping. Cellbender was also used to distinguish nuclei from empty droplets. Background RNA was removed and filtered feature-barcode matrices from Cellbender output were used for downstream primary analyses using scFlow pipeline^89^. Cellbender feature-barcode matrices from a previous sequencing run^22^ (n=19) were integrated with the current cohort for processing with scFlow. Cells were filtered for ≥500 and ≤20000 total counts and ≥400 and ≤8000 total expressive features, where expressivity was defined as a minimum of 1 count in at least 3 cells. The maximum percentage of counts mapping to mitochondrial genes was set to ≤10%. Doublets were identified using the DoubletFinder algorithm^90^, with a doublets-per-thousand-cells increment of 8 (recommended by 10X Genomics), a pK value of 0.02, and embeddings were generated using the first 10 principal components calculated from the top 2000 most highly variable genes.

### Integration, clustering, and visualisation of snRNA sequencing data

The linked inference of genomic experimental relationships (LIGER)^91^ package was used to calculate integrative factors across samples. LIGER parameters included: k = 30, lambda = 5.0, thresh = 0.0001, max_iters = 100, knn_k = 20, min_cells = 2, quantiles = 50, resolution = 1, num_genes = 3000, centre = false. Two-dimensional embeddings of the LIGER integrated factors were calculated using the uniform-manifold approximation and projection (UMAP) algorithm^92^ with the following parameters: pca_dims = 50, n_neighbours = 35, init = spectral, metric = Euclidean, n_epochs = 200, learning_rate = 1, min_dist = 0.3, spread = 0.85, set_op_mix_ratio = 1, local_connectivity = 1, repulsion_strength = 1, negative_sample_rate = 5, fast_sgd = false. The Leiden community detection algorithm was used to detect clusters of nuclei from the UMAP (LIGER) embeddings; a resolution parameter of 0.001 and a k value of 50 were used^93^.

### Initial assignment of cell type labels to snRNA-seq nuclei

After the initial clustering step, automated cell-typing was performed as previously described using the Expression Weighted Celltype Enrichment (EWCE) algorithm^94^ in scFlow^89^, sampling 10,000 cells, against a previously generated cell-type data reference from the Allen Human Brain Atlas^95^. The top five marker genes for each automatically annotated cell-type were determined using Monocle3^96^ and validated against canonical cell-type markers.

### Re-clustering of the snRNA sequencing neuronal subpopulation

In R (v4.4.3; Supplementary Table 21), excitatory and inhibitory neuronal clusters were first subset from the single cell object generated in the initial clustering and cell-type annotation. Data were normalised and scaled using Seurat’s NormalizeData and ScaleData functions. RunPCA was used to compute principal components using the top 2000 highly variable genes. The number of significant PCs was determined empirically: the first PC at which the cumulative variance explained exceeded 90% and the individual variance contribution fell below 5% (co1), and the PC marking the last appreciable drop in variance (>0.1% between consecutive PCs; co2) were calculated, and the minimum of these two values was used. Individual samples were integrated with Harmony^97^, using Seurat’s RunHarmony() function (group.by.vars = “manifest”). To produce the first subcluster UMAP, we used the following parameters in RunUMAP(): dims = 1:16, n.epochs = 200. To identify clusters, we used first the function FindNeighbors() (dims = 1:16) and performed unbiased clustering with FindClusters() (resolution = 0.5). Automated cell-typing was performed essentially as previously described using the EWCE algorithm in scFlow against a previously generated cell-type data reference from the Allen Human Brain Atlas.

For the second round of subclustering, non-neuronal clusters identified in the first clustering round were removed. The existing Harmony embedding was retained, and UMAP (RunUMAP(): dims = 1:17, n.epochs = 200), neighbourhood graph reconstruction (FindNeighbors(): dims = 1:17), and clustering (FindClusters(): resolution = 0.4) were re-run on the filtered neuronal population. EWCE was used to ensure the absence of non-neuronal clusters, resulting in the exclusion of one cluster identified as oligodendrocyte precursor-like before marker gene identification. Neuronal subclusters were annotated based on the differential expression of previously reported canonical entorhinal neuronal subtype-defining genes, including layer-specific excitatory markers (e.g., *CUX2*, *RORB*, *TLE4*, *PCP4*, *THEMIS*, *RELN*, *BCL11B*) and inhibitory neuron markers (e.g., *PVALB*, *SST*, *VIP*, *LAMP5*, *NPY*), primarily as reported previously^23^. Subcluster labels encode broad class (Exc/Inh), cortical layer where applicable, and the top distinguishing marker genes (e.g., Exc-L2-3-*CUX2*-*CALB1*, Inh-*VIP*-*RELN*).

### Dirichlet modelling of relative snRNA sequencing neuronal cell type composition

Per-sample neuronal subcluster proportions were computed using a custom frequency extraction function applied to cell-level metadata. Proportions were transformed using the DR_data() function from the DirichletReg^98^ R package and modelled using Dirichlet multinomial regression (DirichReg()), which accounts for the compositional, bounded nature of cell-type proportions. The primary model was fitted with diagnosis (Control vs AD; Control as reference) as the independent variable of interest, and sex, age, and study cohort as covariates. *APOE* genotype was not applied as a covariate due to the genotype being unknown for 9 of the 36 samples analysed. P values were corrected for multiple comparisons across neuronal subclusters using the Benjamini-Hochberg method, and significance was defined as adjusted p < 0.05. Subclusters with a statistically significant negative coefficient for diagnosis (i.e., reduced proportion in AD relative to controls) were classified as vulnerable; all remaining subclusters were classified as resilient.

### Neuronal subcluster marker gene identification and cross-dataset comparison

Neuronal subcluster marker genes were identified using the FindAllMarkers function from Seurat, run separately within excitatory and inhibitory neuronal subsets. Only positively enriched markers were considered (only.pos = TRUE), with genes required to be detected in at least 25% of cells within the cluster (min.pct = 0.25) and to show a minimum average log2 fold-change of 0.25 relative to all other neurons within the same broad class (logfc.threshold = 0.25). Differential expression was assessed using the Wilcoxon rank-sum test with Bonferroni correction for multiple testing, consistent with the approach used in a previous report^12^.

To assess the transcriptomic correspondence between our neuronal subclusters and those reported in a previous publication investigating selective neuronal vulnerability in AD^12^, pairwise Jaccard similarity was computed between the top 100 marker genes (ranked by average log2 fold-change; adjusted p < 0.05) of each excitatory subcluster in our dataset and each excitatory subcluster in the published atlas, using the marker gene lists provided. Non-cortical clusters (DG granule cells, CA1 pyramidal cells, CA2/CA3 pyramidal cells, and Exc *NXPH RNF220*) were excluded from the published reference prior to comparison. Jaccard similarity was defined as the size of the intersection divided by the size of the union of the two marker gene sets. The resulting similarity matrix was visualised as a heatmap.

### Marker gene-based label transfer and Dirichlet compositional analysis

Previously reported cortical excitatory cluster signatures^12^ (27 clusters retained after excluding 5 clusters absent from the EC) were built from each cluster’s top 25 cluster-discriminating marker genes (adjusted p < 0.05, ranked by avg_log2FC; markers significant in > 3 reference clusters were excluded; restricted to genes in our data). AUCell scores (aucMaxRank = 5%) on log-normalised counts were z-scored per signature across cells, and each of our excitatory cells was assigned to its top-scoring cluster from the atlas; assignments with a best-to-second-best z-score ratio ≥ 1.20 were retained as confident.

Per-donor proportions of confidently labelled cells across the retained clusters (clusters with <3 cells in <15 donors excluded) were fit with DirichletReg (proportion ∼ Braak + sex + age + study ID). Four pre-specified clusters, based on previously reported depletion with AD (Exc TOX3 TTC6, Exc AGBL1 GPC5, Exc RELN GPC5, Exc RELN COL5A2) were tested with Benjamini-Hochberg correction.

### Differential gene expression

Differential gene expression analysis was performed with a pseudocell strategy coupled with mixed linear models as previously described^99^. Pseudocells were constructed by aggregating the raw UMI count of, on average, 30 nuclei per donor and cell type. For the vulnerability contrast, pseudocells were constructed per neuronal subcluster per donor prior to pooling, to preserve subcluster-level resolution. We used the Limma Trend approach^100^ to identify differentially expressed genes within each cell type with age, sex, *post-mortem* interval, Study ID, scaled pseudocell size, log_2_(pseudocell MT%), and log_2_(pseudocell nUMI) as covariates and donor as a random effect. To assess differential gene expression with AD, we stratified our data by cell type and assigned diagnosis as the independent variable. For the comparison of vulnerable vs resilient neurons, we grouped excitatory neurons and assigned the dependent variable as the metadata column defining whether each neuronal cell type was significantly lost with AD or not. P values were adjusted using the Benjamini-Hochberg method, and differentially expressed genes were defined at adjusted p < 0.05.

### Pathway enrichment

Pathway enrichment analysis was performed using the enrichR (v3.4) package^101^ with the over-representation analysis (ORA) method against the ‘GO_Molecular_Function_2025’, ‘GO_Cellular_Component_2025’, ‘GO_Biological_Process_2025’, and ‘KEGG_2021_Human’ databases. The background gene universe was constructed from the union of all genes detected across the full set of differential expression output files for the analysis, representing all genes expressed in neuronal nuclei. Upregulated (logFC > 0) and downregulated (logFC < 0) differentially expressed genes (padj < 0.05) were submitted separately; gene set size was restricted to between 10 and 250 genes. The false-discovery rate (FDR) was calculated using the Benjamini-Hochberg method, and a significance threshold of FDR ≤ 0.05 and a minimum overlap of 5 genes were applied to restrict visualised pathways to those with robust effect sizes. For figure display, pathways were manually curated from those passing significance threshold; all statistically significant genes and pathways are provided (Supplementary Table 9, 10).

### Tau oligomerisation vulnerability scoring

Directional tau oligomerisation gene sets were derived from the genome-wide CRISPR screen previously reported^28^, in which knockout of individual genes in iPSC-derived neurons was assessed for effects on tau oligomerisation. Genes were classified as tau suppressors (ε > 0; genes whose knockout increases tau oligomerisation, indicating a protective role when present) or tau enhancers (ε < 0; genes whose knockout decreases tau oligomerisation, indicating a vulnerability-associated role when present), filtered by nominal significance (p < 0.05). Gene sets were intersected with genes detected in the snRNA-seq dataset prior to scoring.

AUCell scores were computed for each gene set independently using the AUCell R package^29^, with rankings built from log-normalised counts and an aucMaxRank parameter set to the top 15% of expressed genes per cell. A composite tau vulnerability index was calculated per non-diseased nucleus as the difference between the tau enhancer AUCell score and the tau suppressor AUCell score, such that higher values indicate greater inferred susceptibility to tau oligomerisation. This difference score, along with the individual component scores, was then z-transformed across all neurons prior to visualisation and statistical testing.

Subcluster-level differences in tau vulnerability score were assessed using one-versus-all linear mixed-effects models fitted independently for each neuronal subcluster, using the lmerTest R package. The model formula was: score ∼ is_target_subcluster + Age + Sex + (1|CaseID), where is_target_subcluster is a binary indicator for the subcluster under test. Scoring and statistical testing were performed on nuclei from non-diseased donors only. P values were adjusted for multiple comparisons across subclusters using the Benjamini-Hochberg method.

### Cluster ranking robustness

To test whether the snRNA-seq subcluster vulnerability ranking depends on the specific tau modifier genes, we generated 1,000 expression-decile-matched random gene-set pairs. For each enhancer (n = 563) and suppressor (n = 293) gene, one replacement was drawn from the same decile of mean non-diseased control neuron expression. The deployed workflow was run individually for each pair, and 17 subclusters were ranked by sham vulnerability score.

### Cross-modality cluster mapping

Correspondence between snRNA-seq subclusters and IMC clusters (Supplementary Fig. 7a) was established manually based on expression of defining lineage markers because the sparsity of the IMC panel (16 neuronal subtype markers) compared to the snRNA-seq gene universe (25,454 unique genes) rendered automated cross-modality methods unstable in pilot tests. Where multiple snRNA-seq subclusters expressed a shared IMC marker, such as *CALB1* in Exc-L2-3-*CUX2*-*CALB1* and Exc-L3-*PCP4*-*CALB1*, they were pooled into a single IMC-equivalent label. 5 of 6 inhibitory and 5 of 10 excitatory clusters shared key marker genes across modalities. The unassigned snRNA-seq inhibitory cluster was mapped onto the only remaining unassigned IMC inhibitory cluster, inhibitory cluster 1 (GAD1), which lacked inhibitory subtype marker identity beyond GAD1. All 5 unassigned snRNA-seq excitatory clusters were mapped to IMC excitatory cluster 1 (MAP2), which represented a broad class of excitatory neurons lacking distinctive identity, due to the absence of canonical deep layer excitatory neuron markers in the IMC panel. Of the 5 unassigned snRNA-seq excitatory clusters, 4 had canonical layer 5-6 identity based on marker genes; IMC excitatory cluster 1 neurons were shown to be predominantly situated in layers 5-6 (Fig. 1e).

### Protein-level validation of transcriptionally derived vulnerability to tau oligomerisation

To assess whether transcriptomic vulnerability to tau pathology predicted pathological tau accumulation, cluster-level tau oligomerisation vulnerability scores derived from non-diseased donors were compared with PHF1 signal intensity measured by IMC in AD donors. PHF1 signal intensity was extracted from the arcsinh-transformed IMC expression matrix and restricted to neuronal cells. Tau vulnerability scores were aggregated to IMC neuronal clusters using the manual mapping described above, weighted by RNA cell count where multiple snRNA-seq clusters mapped to a single IMC cluster. For each IMC donor, mean PHF1 signal was computed per neuronal cluster, yielding a pseudobulk profile across 10 clusters.

The Spearman rank correlation between cluster-level tau vulnerability Z-score and mean PHF1 signal was computed independently for each AD donor. Per-donor correlations were Fisher z-transformed and tested against zero using a one-sample t-test (df = number of donors – 1), with the mean correlation and 95% confidence interval back-transformed to the rho scale for reporting. This approach treats the IMC donor as the unit of replication, giving correctly calibrated inference for a cluster-level predictor derived from an independent cohort.

### Correlation of tau oligomerisation vulnerability gene expression with PHF1 immunostaining

To characterise the relationship between expression of individual genes from the CRISPR screen^28^ and tau phosphorylation burden at the neuronal subtype level, mean log-normalised expression of each gene in the tau oligomerisation gene sets was computed per snRNA-seq neuronal subcluster in non-diseased control donors. Subcluster-level expression profiles were mapped to IMC neuronal clusters using a cell-count-weighted aggregation scheme identical to that used for tau vulnerability score regression, in which RNA subclusters mapping to the same IMC cluster were combined as weighted means proportional to subcluster cell count. This yielded a gene expression profile across ten IMC neuronal clusters for each tau oligomerisation vulnerability gene.

For each gene, the Spearman rank correlation between cluster-level expression and mean PHF1 signal intensity was computed independently for each AD donor with all three ROIs passing QC (n = 44). Per-donor Spearman rho values were Fisher z-transformed and tested against a null hypothesis of zero mean correlation using a one-sample t-test, with the mean rho back-transformed to the rho scale for reporting. Multiple testing correction was applied using the Benjamini-Hochberg method both within each gene set (enhancer and suppressor separately) and across all tested genes combined. Genes were classified as concordant where the direction of PHF1 correlation was consistent with their experimentally determined role in the screen (positive rho for enhancers; negative rho for suppressors) and discordant otherwise. Set-level directional enrichment was assessed using one-sample Wilcoxon signed-rank tests against a null of zero median rho, with one-sided alternatives appropriate to each gene set direction.

### Protein-protein interaction network analysis

Protein-protein interaction network analysis was performed on the top 100 concordant genes from each gene set, ranked by magnitude of mean per-donor Spearman rho. Interactions were retrieved from STRING (v12.0, species: *Homo sapiens*) using the STRINGdb R package^102^, with a combined interaction score threshold of 0.7. Evidence was restricted to experimental, database-curated, and co-expression channels; text-mining evidence was excluded. Self-loops and duplicate symmetric edges were removed. Genes with no qualifying interactions within the input gene set were excluded from visualisation. Networks were constructed as undirected graphs using tidygraph and visualised using ggraph with a Fruchterman-Reingold force-directed layout. Node fill was mapped to Spearman rho using a diverging blue-white-red colour scale; node size was mapped to network degree.

### Gene set enrichment analysis

GSEA was performed on concordant genes, those for which the direction of PHF1 correlation was consistent with their experimentally determined effect on tau oligomerisation in the CRISPR screen, ranked by mean per-donor Spearman rho. Gene symbols were converted to Entrez IDs using the bitr function from clusterProfiler^103^ with org.Hs.eg.db as the annotation database; genes failing to map were excluded. Where a symbol mapped to multiple Entrez IDs, the first was retained. GSEA was performed against the Reactome pathway database using gsePathway from ReactomePA^104^, with the following parameters: minimum gene set size 15, maximum gene set size 500, 1000 permutations, Benjamini-Hochberg p-value adjustment, significance threshold p.adjust < 0.05. A fixed random seed (42) was applied prior to all permutation-based analyses to ensure reproducibility. Pairwise term similarity was computed using enrichplot::pairwise_termsim for downstream visualisation. GSEA was additionally performed against GO Biological Process (clusterProfiler::gseGO) and KEGG (clusterProfiler::gseKEGG) databases using identical parameters; no significant terms were identified in either database at the specified threshold and these results are not reported.

### Trajectory analysis

Braak pseudotime trajectory analysis was performed to infer phenotypic transitions happening from early to late Braak stages. Unsupervised single-cell trajectory analysis was performed with Monocle3, an algorithm that learns the sequence of gene expression changes each cell must go through as part of a dynamic biological process. Prior to trajectory analysis, the cluster of interest was re-processed: highly variable genes were identified using variance-stabilising transformation (nfeatures = 3000), data were scaled, and PCA was performed (npcs = 40). SeuratWrappers (v0.4.0) was used to convert the Seurat object into a CellDataSet (cds) object. We pre-processed the cds object with num_dim equal to the minimum of either the first PC which cumulatively explains >90% of the variance and where the individual PC explains <5%, or the PC with the last significant drop in variance, then performed a batch correction to integrate at the sample level with Batchelor^105^ using the align_cds() function with alignment_k = 20. During batch correction, we also regressed out covariates using residual_model_formula_str = “∼ total_features_by_counts + pc_mito + Sex + Age + StudyID + PostMortemInterval”. A dimension reduction was performed using reduce_dimension() with default settings, before we ran cluster_cells() (resolution = 0.001). Following initial clustering, one cluster was identified as a technical artefact driven by a single donor and was excluded. Batch correction, dimensionality reduction, and clustering were subsequently repeated on the dataset. The learn_graph() function (use_partition = FALSE, close_loop = FALSE, learn_graph_control = list(rann.k=65, prune_graph = TRUE, orthogonal_proj_tip = FALSE, minimal_branch_len = 12, ncenter = 135)) was used to learn the trajectory. The origin of the trajectory was assigned manually as the node with the greatest proportion of non-diseased nuclei. To identify genes differentially expressed along the trajectory we used the graph_test() function (neighbor_graph = “principal_graph”, k = 50). Trajectory-associated genes from graph_test() were pre-filtered to those with q ≤ 0.05 and Moran’s I above the 70^th^ percentile, further restricted to genes expressed in ≥ 1% but ≤ 90% of nuclei with mean expression between 1 and 100 normalised counts. This gene set was then passed to find_gene_modules() (resolution = 0.05, umap.metric = “euclidean”, umap.min_dist = 0.3, umap.n_neighbors = 50L, umap.fast_sgd = FALSE). Module gene sets were tested for pathway enrichment using enrichR against GO Biological Process, Molecular Function, Cellular Component (2025), WikiPathways (2023), KEGG (2021), BioPlanet (2019), and the Kinase Library (2023) databases. Terms were retained at FDR ≤ 0.05 with a minimum gene overlap of 3.

### Robustness of pseudotime ordering to root selection

To test whether the Exc-L2-3-*CUX2*-*CALB1* pseudotime ordering depended on root choice, the principal graph fit by monocle3::learn_graph was held fixed and order_cells was re-run with ten different root nodes: the five principal-graph vertices most enriched for nuclei from non-diseased control brains, and the five most enriched for nuclei from AD brains (Supplementary Fig. 8b), ranked by per-vertex enrichment ratio over the cohort baseline (vertices with < 20 assigned cells excluded). Pairwise Spearman correlations between resulting per-cell pseudotime vectors were computed and visualised as a heatmap (Supplementary Fig. 8c). Median within-end correlations were ρ = 0.85 (non-diseased control) and 0.97 (AD), supporting the conclusion that the inferred pseudotime ordering is robust to root selection within each end of the trajectory.

### Connectivity Map analysis

The resilience signature was assembled by filtering the tau-modifier gene set^28^ for those expressed in concordance with cluster level PHF1 antibody signal and intersecting these genes with those differentially expressed in our Exc-L2-3-*CUX2*-*CALB1* contrast (suppressors ∩ downregulated DEGs; enhancers ∩ upregulated). After restricting to the LINCS L1000 gene universe^106^, 29 up-signature (upregulation associated with resilience) and 37 down-signature (downregulation associated with resilience) genes remained.

The signature was queried against LINCS L1000 Phase I profiles via the signatureSearch package^107^ using the Weighted Tau Connectivity Score (WTCS; gess_lincs), restricted to neural cell contexts (NPC, FIBRNPC, NEU). For each compound (pert), the instance with the largest |NCS| was retained, preserving sign. Tool/probe compound prefixes (BRD-, SA-, STK-, CAY[0-9], XMD-, HG-, ALW-, QL-, WYE-, LY[0-9]) were excluded and a WTCS FDR ≤ 0.05 cutoff was applied. MoA annotations were taken from the LINCS MOAss field augmented by the Broad Institute compoundinfo_beta.txt (clue.io), matched on pert_iname then pert_id; multi-MoA strings were expanded across all annotated classes (split on ; or |).

### Blood-brain barrier penetrance annotation

Each compound was scored for predicted CNS suitability using the six-property CNS-MPO^108^ (0-6 scale) from RDKit descriptors (Crippen.MolLogP, Descriptors.MolWt, rdMolDescriptors.CalcTPSA, rdMolDescriptors.CalcNumHBD); cLogD_7.4_ was substituted with cLogP, and basic pKa was estimated via a SMARTS detector of ionizable nitrogenous groups (primary, secondary and tertiary aliphatic amines, guanidines, amidines, aromatic amines, pyridine and piperazine nitrogens) returning the highest matched literature-default pKa; non-ionizable compounds defaulted to the maximal pKa sub-score. Each property was scored piecewise-linearly and summed.

Empirical BBB± labels were taken from the B3DB curated database^109^. To ensure library-consistent matching, InChI keys were recomputed via RDKit for both the LINCS canonical SMILES and the B3DB InChI strings, then joined on the 14-character connectivity-block skeleton. Compounds were classified as BBB+ confirmed (B3DB BBB+, or CNS-MPO ≥ 5 in the absence of empirical data), BBB+ likely (no empirical data, CNS-MPO 4-5), uncertain (no empirical data, CNS-MPO 3-4), or BBB- likely (B3DB BBB-, or CNS-MPO < 3 in the absence of empirical data). When both signals were available, the empirical B3DB call took precedence.

### Drug candidate prioritisation

Translationally prioritised candidates were defined as compounds with NCS > 1, WTCS FDR ≤ 0.05, and BBB+ confirmed or likely. Per-MoA aggregation retained classes containing ≥ 5 such candidates, ranked by median candidate NCS.

## Code availability

Analysis scripts are published on GitHub (https://github.com/slboulger/EC_Neuronal_Vulnerability).

## Data availability

Source data are supplied in Supplementary Tables where applicable and will be made public after peer review. Raw and processed snRNA-seq data produced for this study will be available on the Gene Expression Omnibus (GEO) database (https://www.ncbi.nlm.nih.gov/geo/) after peer review. Raw and processed snRNA-seq data from the previously published dataset^22^ integrated with the novel dataset are available under the ID GSE264648. Raw and processed IMC data will be available on Figshare following peer review.

## Design of figures

Plots were generated in RStudio, and figures were arranged in Microsoft^®^ PowerPoint for Mac (v16.107.4). Fig. 1b was generated in BioRender (Boulger, S. (2026) https://BioRender.com/q1qqafd).

## Acknowledgements

We thank the study participants, their families, and the coordinating brain banks, without whom this study would not have been possible. We thank all members of the Matthews laboratory at the UK Dementia Research Institute at Imperial College London for discussions and feedback. We thank Megan Winterbotham and Eduardo Lopez Tobar for laboratory management and support and Josh Beale for organisational and administrative assistance.

## Author contributions

This study was conceived and directed by A.C. and P.M.M.. Planning and ordering of the human brain samples was managed by S.L.B. and A.C.. The antibody panel for IMC was curated by S.L.B. and A.C., and IMC antibody conjugation, staining, and acquisition was done by S.L.B., D.L., and A.C.. S.L.B. analysed the IMC data. Y.Z., H.K., and P.N. performed validation experiments. Nuclei isolation was performed by S.L.B. with technical support from E.A. and M.P., and sequencing was done by Imperial Genomics Facility. S.L.B. analysed the single nucleus RNA sequencing data using code written by S.L.B., B.A., M.T., N.N.F., and E.P.D.. A.M., E.P.D., A.C., and P.M.M. provided supervision. S.L.B. drafted the manuscript with P.M.M.. All authors reviewed and edited the manuscript.

## Supplementary Figures

**Supplementary Fig. 1.**
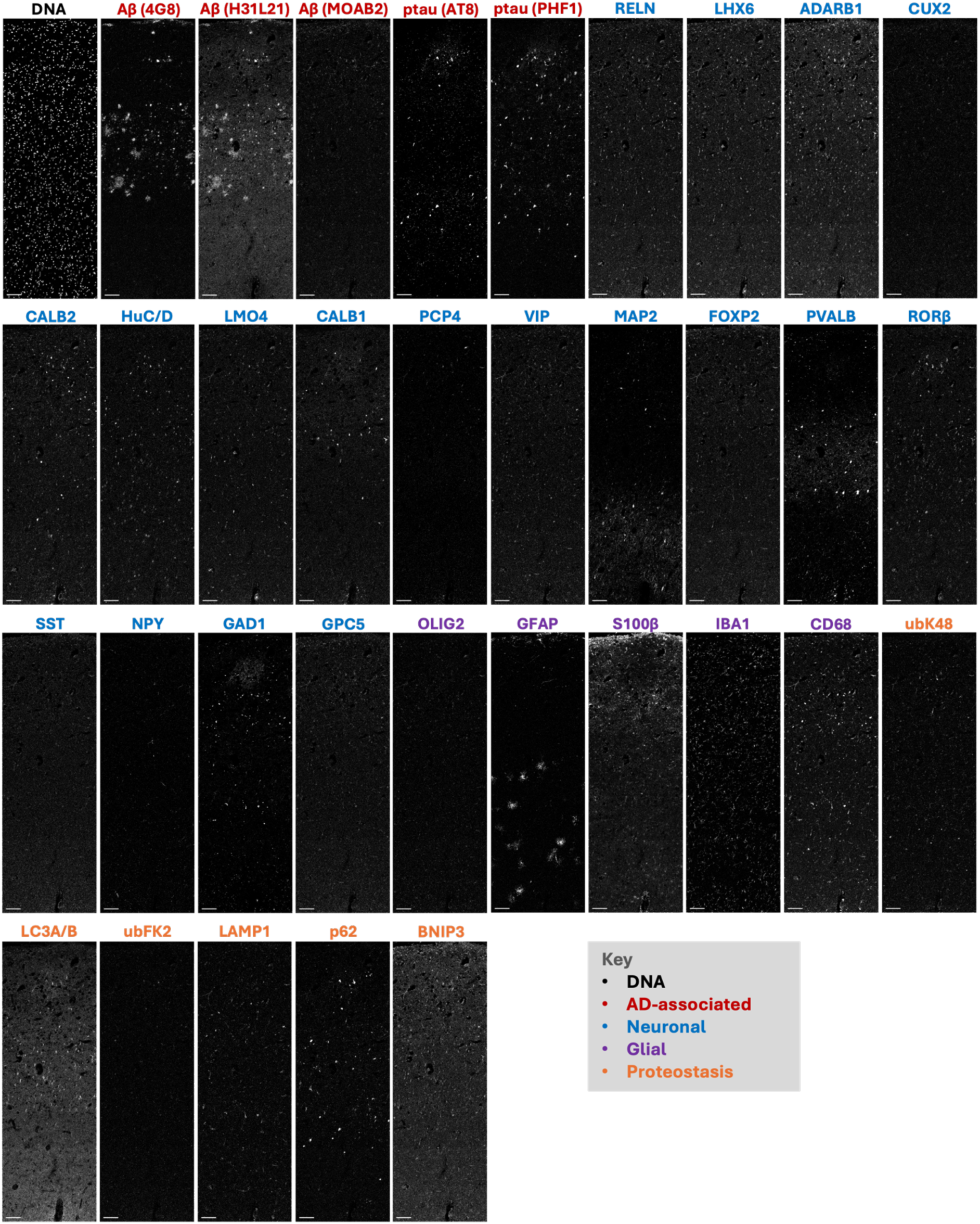
Full imaging mass cytometry antibody panel applied to post-mortem human entorhinal cortex. Representative single-channel IMC images from a single region of interest (ROI), showing signal intensity of each of the 34 protein markers and a DNA intercalator in the panel. Images were generated using Steinbock^80^ and span the full grey matter depth, oriented from the white matter (bottom) to the pial surface (top); scale bar = 100um. Note that OLIG2 was excluded from downstream analysis due to suboptimal staining performance.

**Supplementary Fig. 2.**
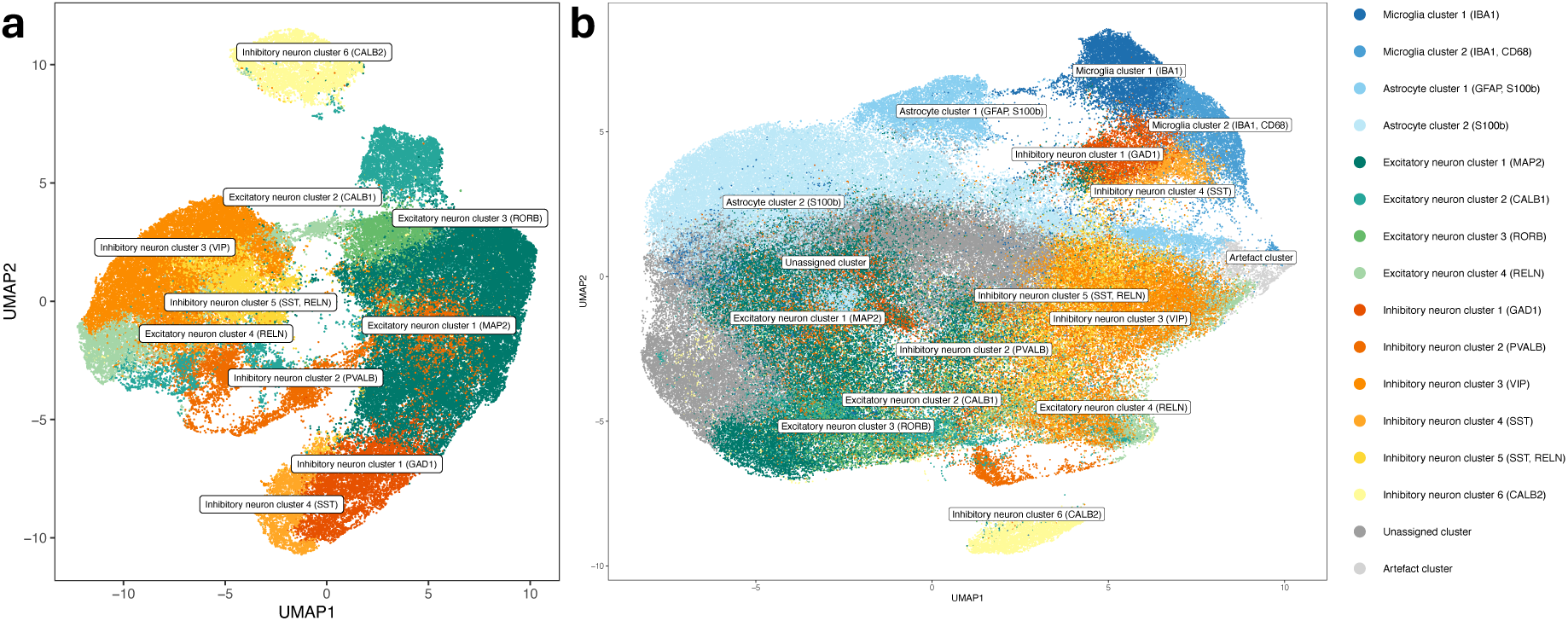
Two-stage IMC cell phenotyping strategy resolving neuronal subpopulations across the entorhinal cortex. **a**, UMAP embedding of neurons after the first phenotyping stage. PhenoGraph clustering identified 17 neuronal subclusters, which were subsequently annotated and merged based on differential marker abundance into 4 excitatory and 6 inhibitory neuronal populations. **b**, Full IMC UMAP, with neuronal subpopulation labels from **a** projected back onto the original UMAP containing neuronal and non-neuronal cells. Colours correspond to cell type identities as shown in the legend.

**Supplementary Fig. 3.**
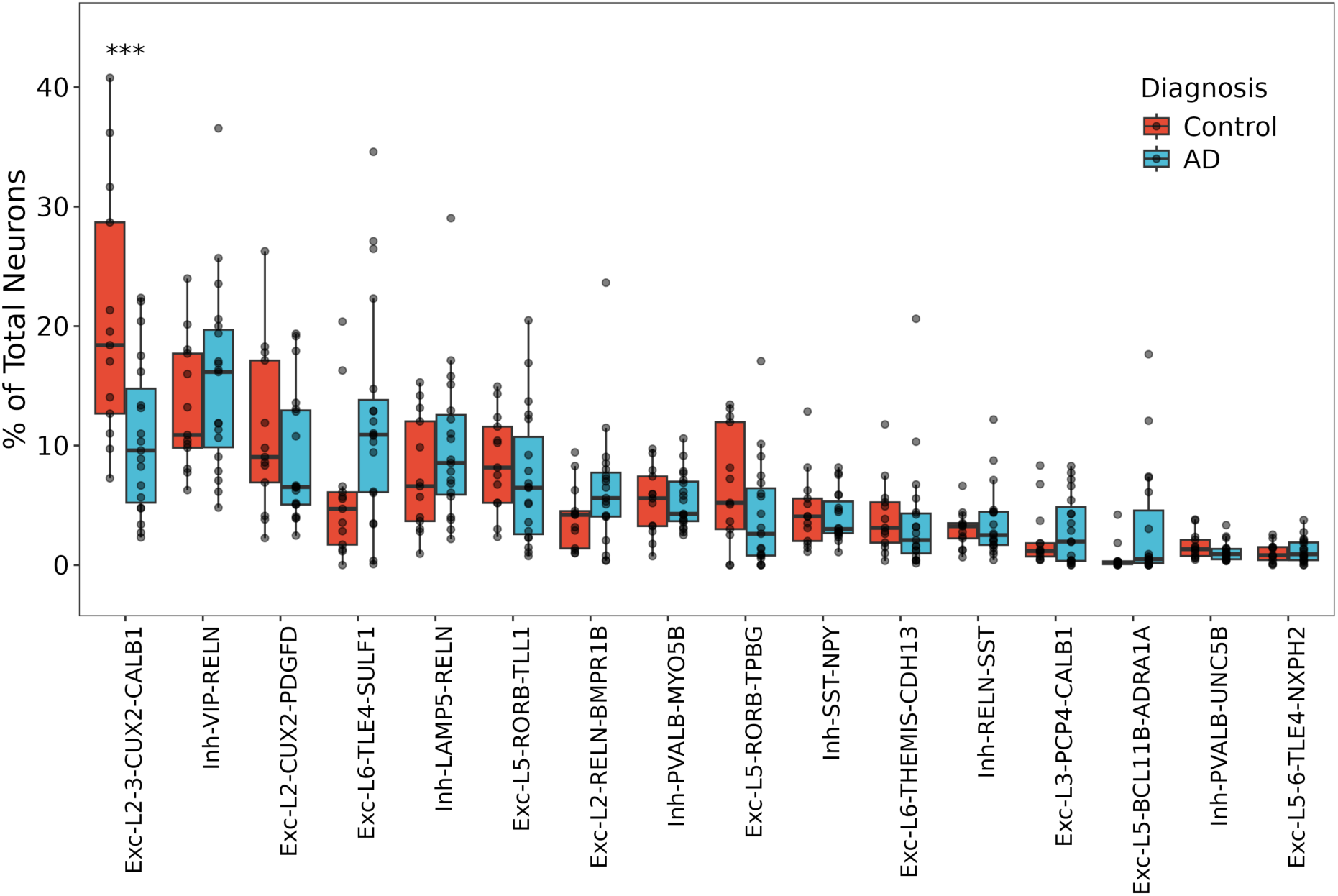
Neuronal snRNA-seq subcluster abundance with the exclusion of samples with early AD-related pathology (Braak 2). Proportional abundance of snRNA-seq neuronal populations in AD (Braak 3-6) versus control (Braak 0-1) donors, ordered by population size. Statistics derived from Dirichlet multinomial regression (proportion ∼ diagnosis + age + sex + study ID). n = 13 control donors, 19 AD; snRNA-seq cohort. Each point represents one donor. *** p < 0.001.

**Supplementary Fig. 4.**
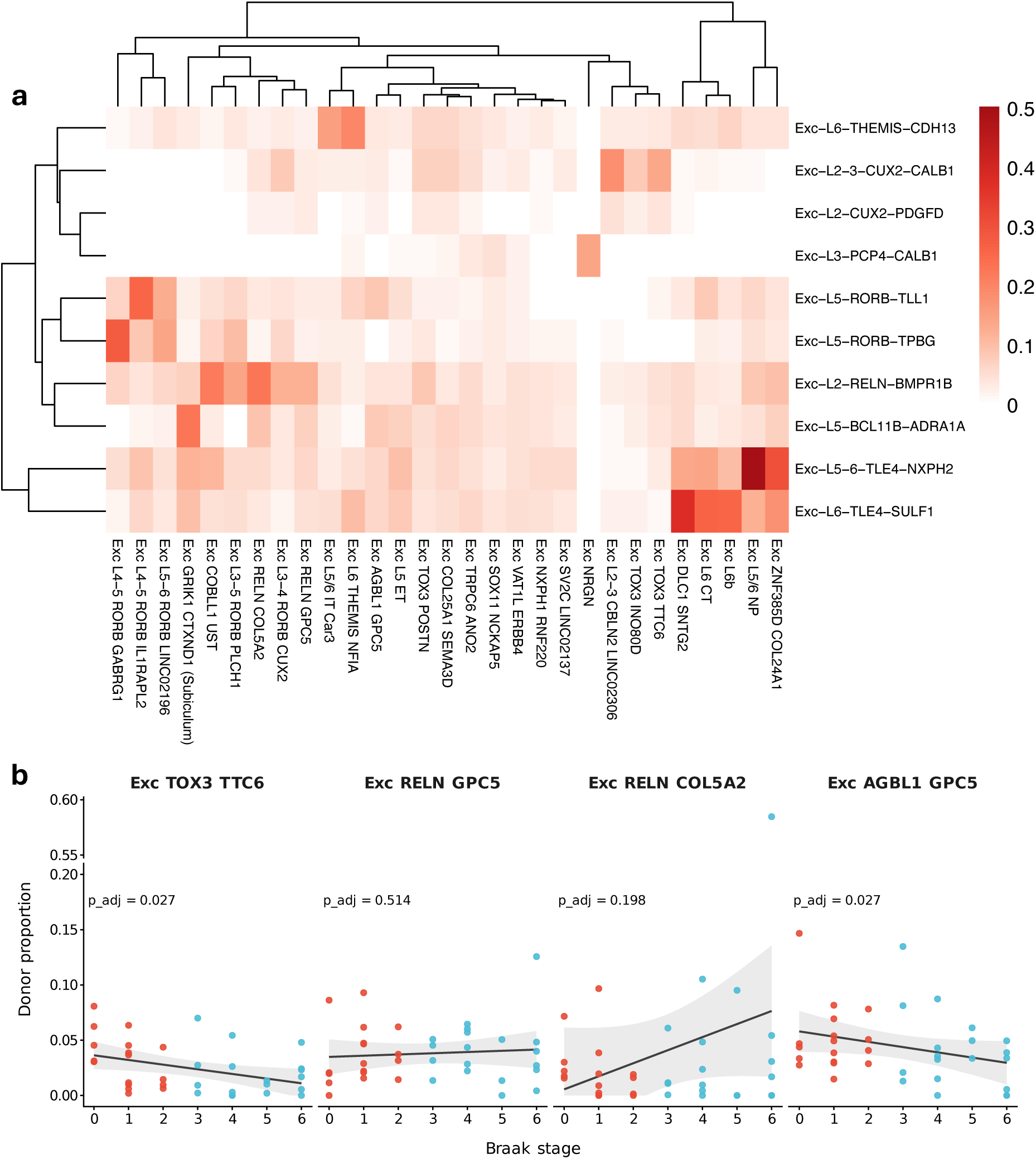
**Transcriptomic similarity between excitatory neuron subclusters from the present study and a previously published atlas**^12^**. a,** Heatmap showing pairwise Jaccard similarity coefficients computed from the top 100 marker genes (ranked by average log2 fold-change) for each excitatory neuron subcluster identified in the present study (y-axis) and EC-associated excitatory neuron clusters from a previously published study^12^ (x-axis). Hippocampal and thalamic clusters not represented in the EC were excluded from the reference dataset. Colour scale indicates Jaccard similarity from 0 (white) to maximum observed similarity (red). Rows and columns are ordered by hierarchical clustering. **b,** Per-donor proportions of confidently labelled cells for four neuronal subclusters reported to be depleted previously following marker-based label transfer (AUCell against the top 25 cluster-discriminating marker genes; per-cluster z-scoring; confident assignment = ratio of best-to-second-best z-score ≥ 1.20; n = 36 donors). Points coloured by clinical diagnosis: red, non-diseased; blue, AD. Grey lines show ordinary least squares fit with 95% confidence interval for visualisation. P-values are BH-adjusted from a Dirichlet model (proportion ∼ Braak + sex + age + study ID).

**Supplementary Fig. 5.**
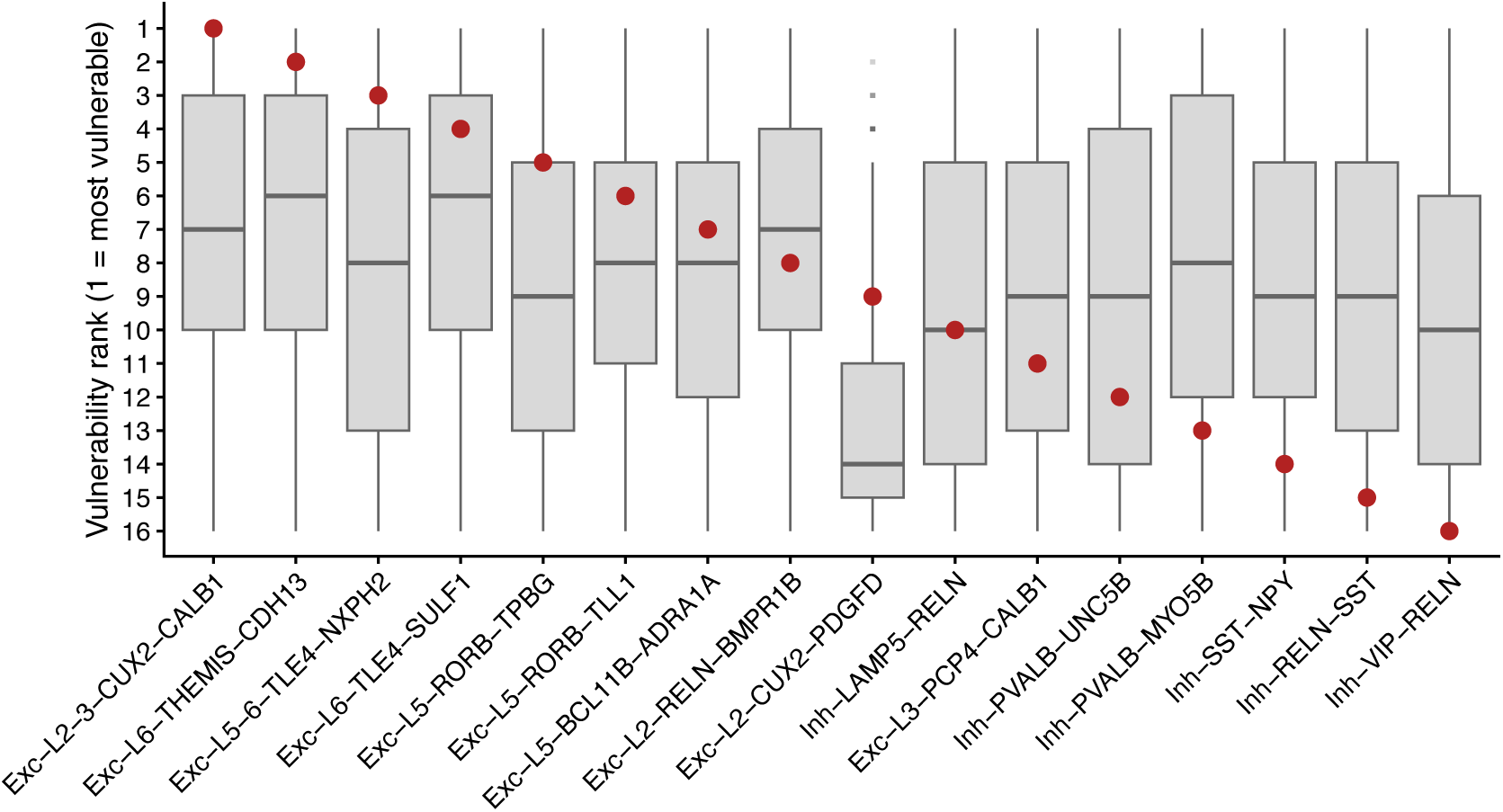
Subcluster vulnerability rank under expression-matched random gene sets and with exclusion of samples with early AD-related pathology (Braak 2). a, Per-iteration rank distribution for each of the 17 snRNA-seq neuronal clusters across 1000 random expression-decile-matched enhancer (n = 563) and suppressor (n = 293) gene-set pairs, scored via the deployed tau oligomerisation scoring workflow. Box plots show the median, IQR, 1.5 x IQR, and outliers. Red dots are ranks obtained with the actual tau oligomerisation gene sets.

**Supplementary Fig. 6.**
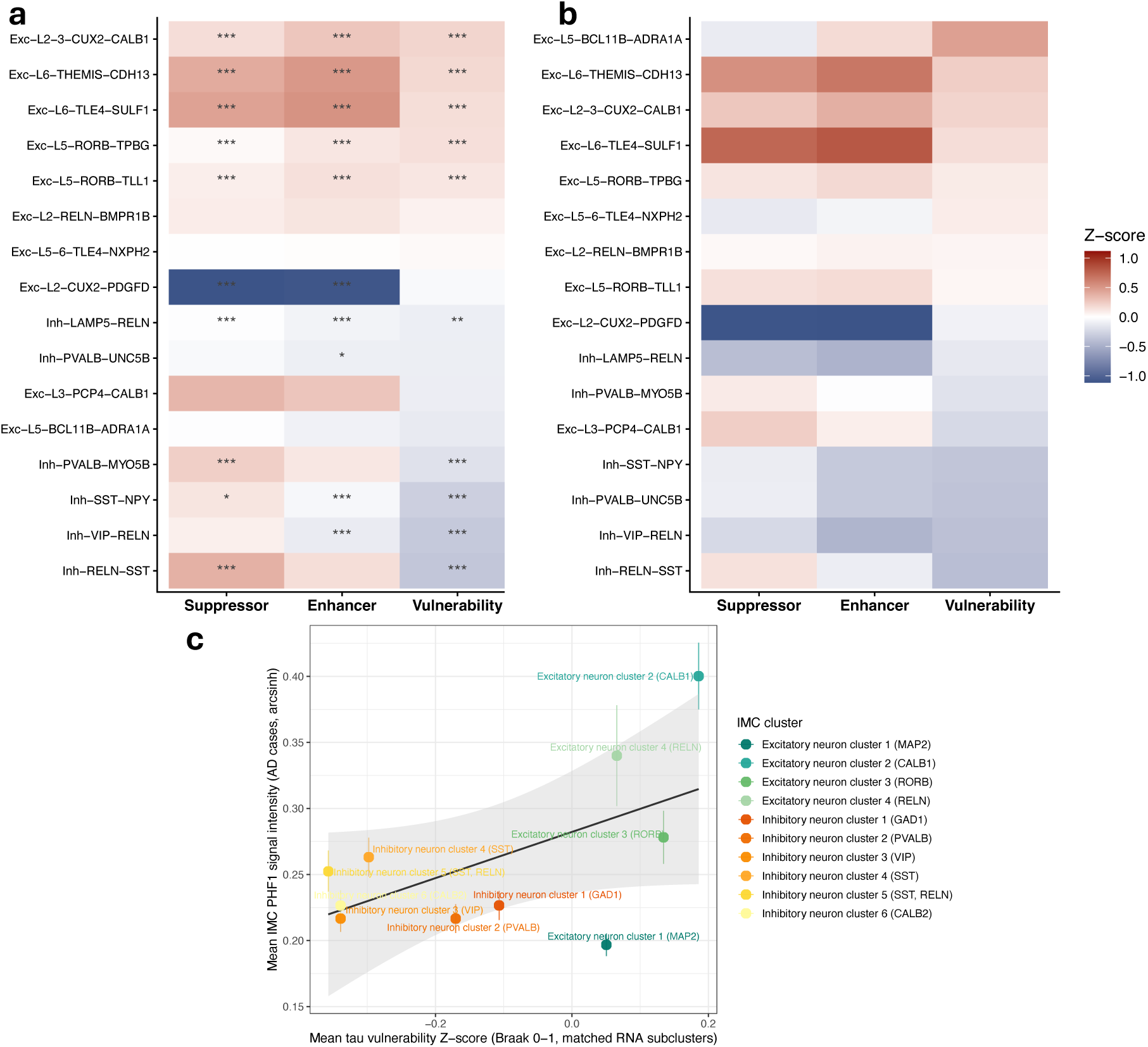
Subcluster tau oligomerisation vulnerability with exclusion of samples with early AD-related pathology (Braak 2). a-b, AUCell-derived scores for tau oligomerisation enhancer and suppressor gene sets and a composite vulnerability score (enhancer minus suppressor activity) across neuronal subclusters in Braak 0-1 **(a)** and Braak 0 only donors **(b)**. Heatmap displays z-scored module activity per subcluster. Subcluster differences were tested using one-versus-all linear mixed-effects models (score ∼ subcluster + age + sex + (1|donor)); p values were adjusted using the Benjamini-Hochberg method. *FDR < 0.05, **FDR < 0.01, ***FDR < 0.001. n = 5 Braak 0 donors and 8 Braak 1 donors; snRNA-seq cohort. **c,** Mean PHF1 signal intensity per IMC neuronal cluster plotted against tau vulnerability Z-score derived from Braak 0-1 donor snRNA-seq. Each point represents a neuronal cluster (n = 10 clusters); error bars indicate SEM across AD donors. The regression line is fit to cluster means for visualisation only and does not reflect the inferential analysis (see text). Tau vulnerability scores and PHF1 measurements were obtained from independent cohorts (non-diseased controls and AD donors, respectively). Statistical inference was performed at the donor level: the Spearman rank correlation between cluster-level tau vulnerability Z-score and mean PHF1 was computed independently for each AD donor and per-donor correlations were Fisher z-transformed and tested against zero using a one-sample t-test (mean ρ = 0.345, 95% CI [0.255, 0.428]; t(43) = 7.35, p = 3.97 × 10⁻^9^; n = 44 AD donors).

**Supplementary Fig. 7.**
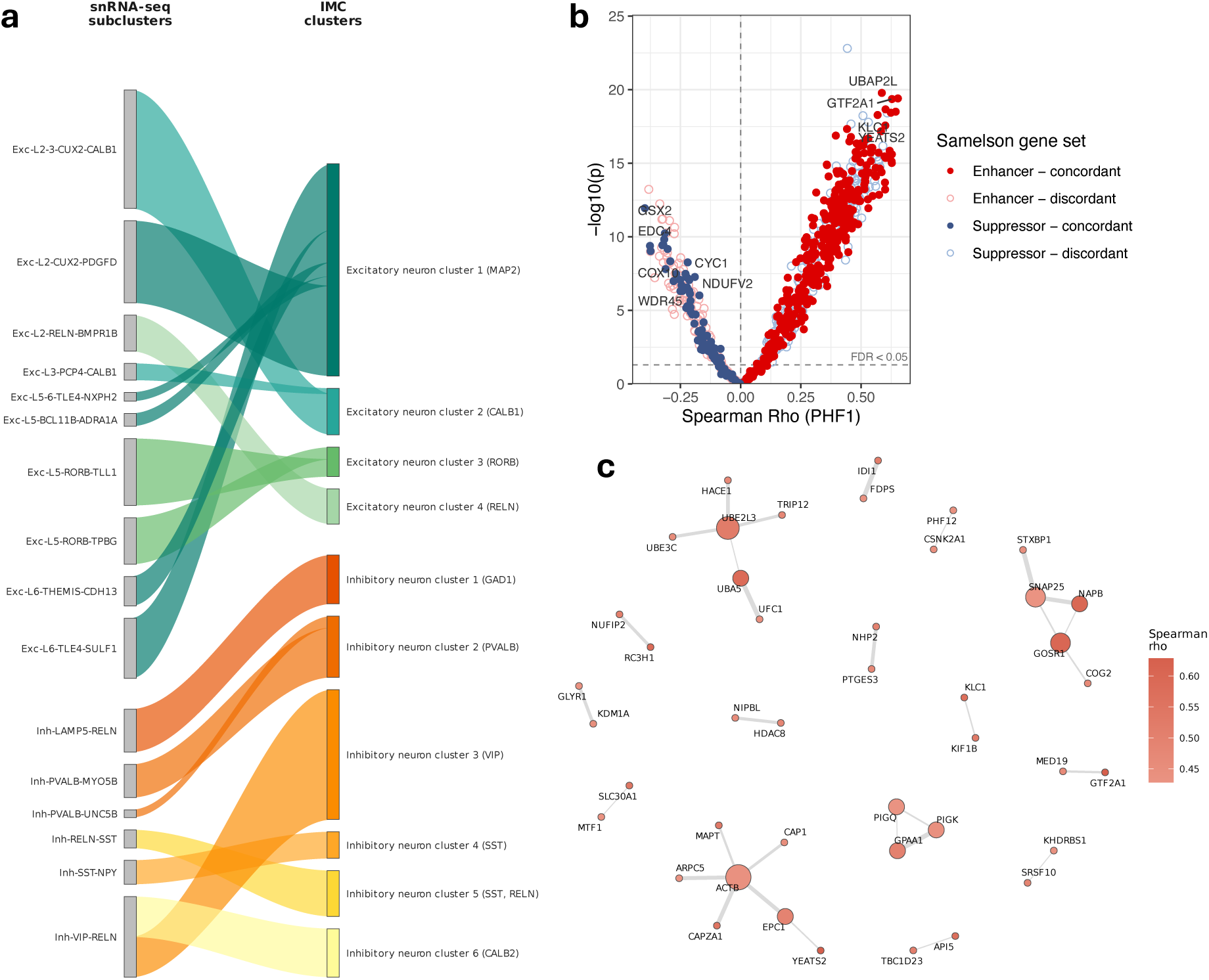
Transcriptomic correlates of tau phosphorylation burden in reported CRISPR screen gene sets. a, Alluvial diagram depicting the manual curation used to establish correspondence between snRNA-seq neuronal subclusters (left) and IMC neuronal clusters (right) for the tau vulnerability score–PHF1 regression analysis. Node width is proportional to the relative abundance of neurons within each cluster. Flows represent assigned correspondences between modalities, coloured by IMC cluster identity. Where multiple snRNA-seq subclusters mapped to a single IMC cluster, tau vulnerability scores were averaged weighted by RNA cell count prior to regression. Excitatory clusters are shown in teal/green and inhibitory clusters in orange/yellow, consistent with cell-type colour conventions used throughout. **b,** Volcano plot of mean per-donor Spearman rank correlations between tau oligomerisation vulnerability gene expression and PHF1 immunoreactivity (pS396/pS404) across IMC neuronal clusters in AD donors (n = 44 AD donors; 10 IMC neuronal clusters). Each point represents one gene from the CRISPR screen, coloured by experimentally determined role in tau oligomerisation (enhancer or suppressor) and by concordance between screen direction and PHF1 correlation with the direction of relative expression with increasing PHF1 burden in the human EC. Filled symbols indicate concordant genes; open symbols indicate discordant genes. The dashed horizontal line indicates FDR < 0.05. Selected genes discussed in the main text are labelled. **c,** STRING protein-protein interaction network of the top 100 concordant tau pathology enhancer genes, ranked by mean per-donor Spearman correlation with PHF1. Nodes are sized by network degree and coloured by Spearman rho.

**Supplementary Fig. 8.**
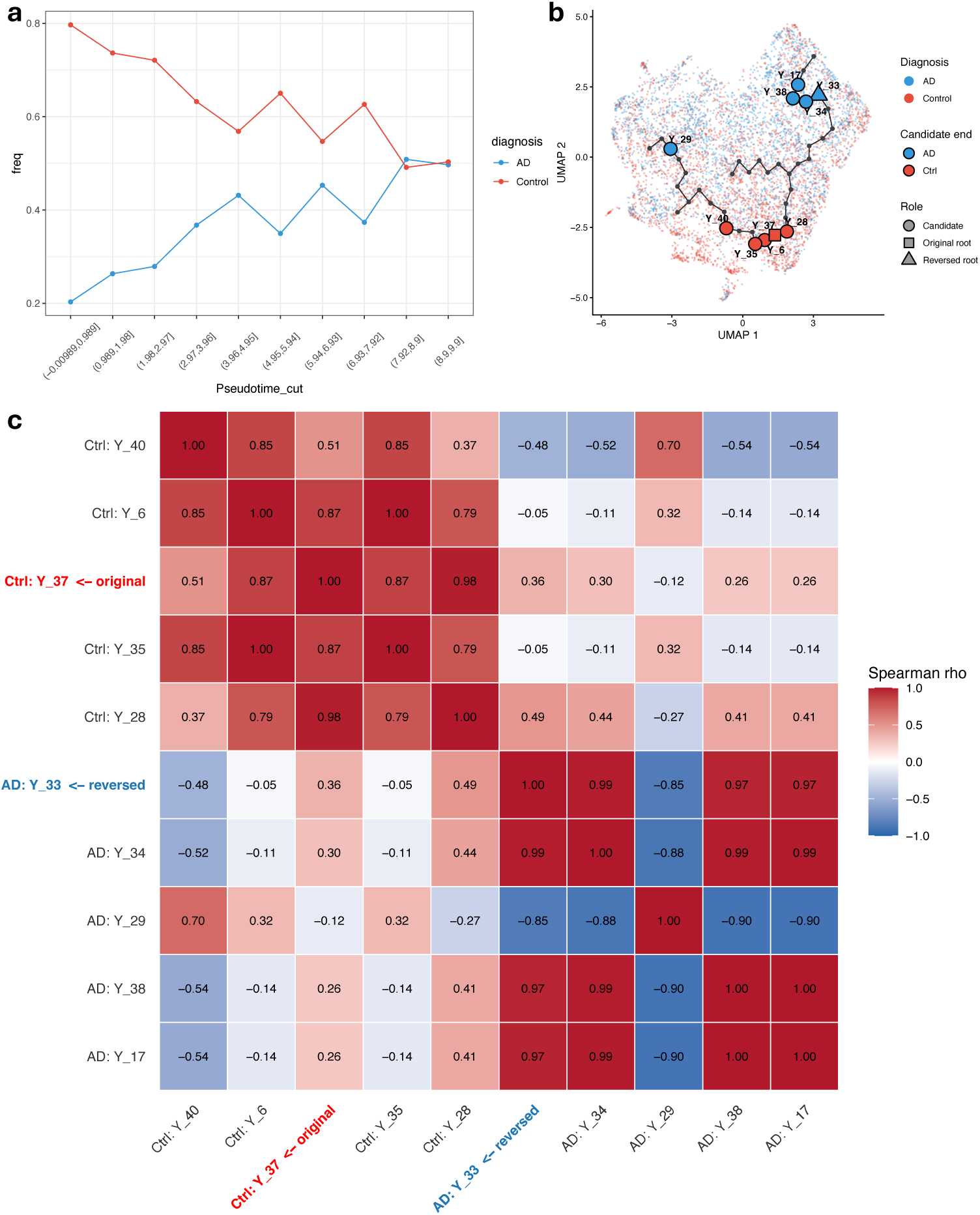
Pseudotime trajectory characterisation and root robustness for the Exc-L2-3-CUX2-CALB1 neuronal population. a, Diagnostic composition along pseudotime. Cells along the Exc-L2-3-CUX2-CALB1 trajectory (Fig. 5e) were partitioned into 10 equal pseudotime bins; the fraction of non-diseased control and AD nuclei per bin is shown. **b,** Ten alternative root nodes are highlighted: the five principal-graph nodes most enriched for control nuclei (red fill; including the published root Y_37, square) and the five most enriched for AD nuclei (blue fill; most-enriched Y_33, triangle). Background cells are coloured by diagnosis (pink, control; light blue, AD). **c,** Pairwise Spearman correlation between cell-level pseudotime orderings under each of the ten candidate root nodes shown in **b**. For each candidate, order_cells was re-run with that node as root_pr_nodes (principal graph and all upstream steps held fixed); the heatmap shows ρ between the resulting per-cell pseudotime vectors. Rows and columns are blocked by candidate end: Control-end candidates (top-left 5×5), AD-end candidates (bottom-right 5×5). The published root Y_37 (red, bold) and the most AD-enriched root Y_33 (blue, bold) are highlighted. Within-end correlations are uniformly high (median ρ = 0.85 among Control candidates, 0.97 among AD candidates), demonstrating that the inferred pseudotime is robust to root selection within each end. Lower within-end pairs involving Y_40 (Control) and Y_29 (AD) reflect these candidates lying on minor side-branches of the principal graph rather than along the main Y_37 to Y_33 axis. n = 17 control donors, 19 AD donors; snRNA-seq cohort.

**Supplementary Fig. 9.**
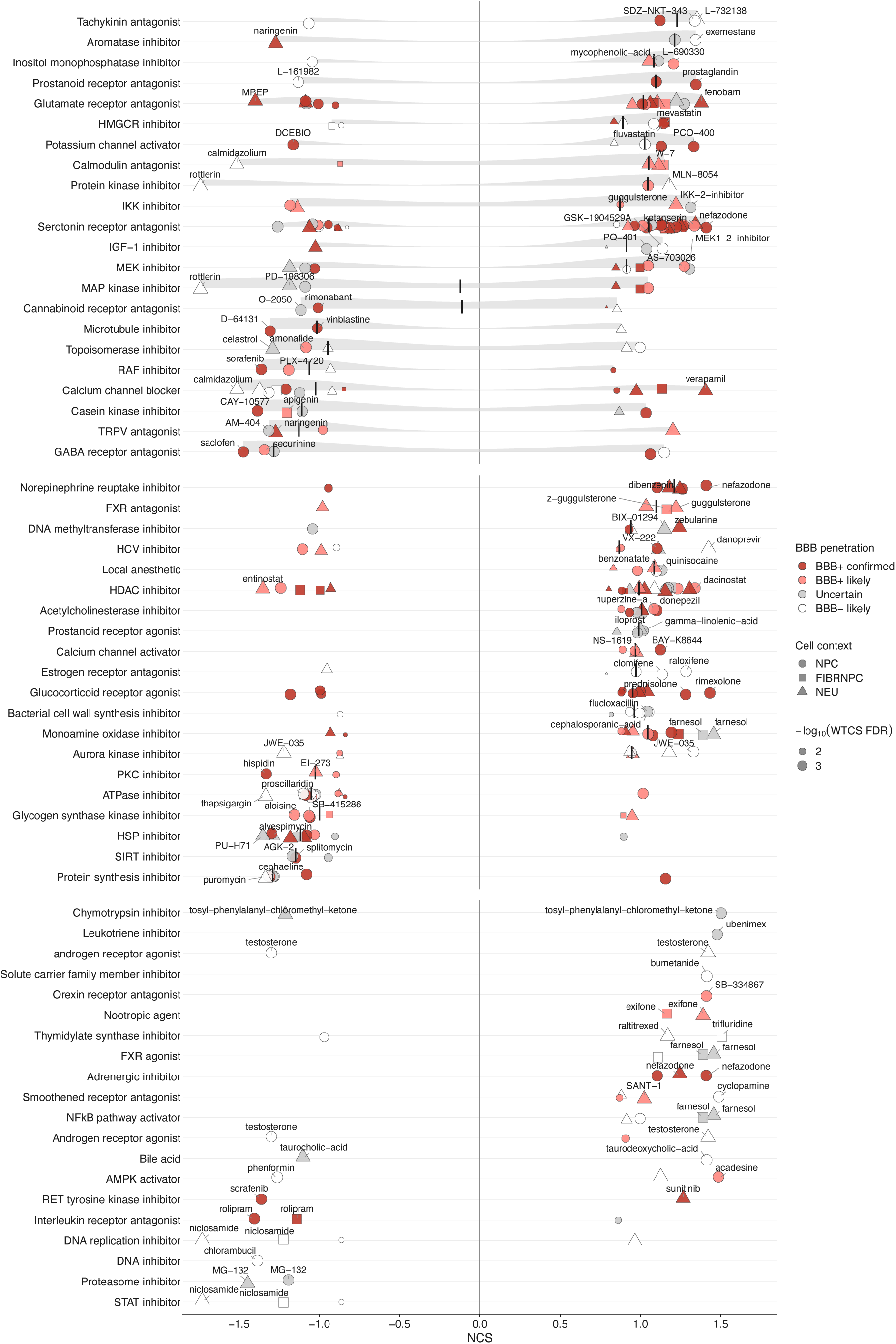
Per-mechanism distribution of compound connectivity scores in the in silico screen of EC tau-pathology resilience modulators. Compound-level NCSs for neural LINCS instances (NPC, FIBRNPC, NEU) passing WTCS BH-adjusted FDR < 0.05. Each y-axis row is one MoA class; dots represent individual (compound × neural cell line) instances at their signed NCS. Compounds are coloured by predicted blood-brain barrier penetration class, combining empirical B3DB labels with the CNS-multiparameter optimisation (MPO) score^108^. Empirical B3DB calls override predicted MPO The short black vertical mark indicates the within-row median NCS where present. MoA rows are partitioned into three groups by overlaid geometry: rows with a half-violin density slab above the points contain MoAs with n ≥ 3 compounds and per-compound NCS interquartile range ≥ 0.5 (broad within-MoA effects, top 2 |NCS| compounds labelled); rows with a median tick but no slab contain MoAs with n ≥ 3 compounds and IQR < 0.5 (concordant within-MoA effects, top 2 labelled); rows with neither slab nor tick are sparse MoAs with ≤ 2 compounds (both labelled). Within groups, rows are ordered by median NCS (mimickers at top, reversers at bottom). The top 20 MoAs per group by median |NCS| are shown, augmented by **Fig. 6c**, yielding 62 classes in total. Tool/probe compounds are excluded; multi MoA compounds contribute one row per tag.

